# TransHLA: A Hybrid Transformer Model for HLA-Presented Epitope Detection

**DOI:** 10.1101/2025.01.20.634002

**Authors:** Tianchi Lu, Xueying Wang, Wan Nie, Miaozhe Huo, Shuaicheng Li

**Author notes:** Contributed equally (Tianchi Lu, Xueying Wang).

## Abstract

**Background:** Precise prediction of epitope presentation on human leukocyte antigen (HLA) molecules is crucial for advancing vaccine development and immunotherapy. Conventional HLA-peptide binding affinity prediction tools often focus on specific alleles and lack a universal approach for comprehensive HLA site analysis. This limitation hinders efficient filtering of invalid peptide segments.

**Results:** We introduce TransHLA, a pioneering tool designed for epitope prediction across all HLA alleles, integrating Transformer and Residue CNN architectures. TransHLA utilizes the ESM2 large language model for sequence and structure embeddings, achieving high predictive accuracy. For HLA class I, it reaches an accuracy of 84.72% and an AUC of 91.95% on IEDB test data. For HLA class II, it achieves 79.94% accuracy and an AUC of 88.14%. Our case studies using datasets like CEDAR and VDJdb demonstrate that TransHLA surpasses existing models in specificity and sensitivity for identifying immunogenic epitopes and neoepitopes.

**Conclusions:** TransHLA significantly enhances vaccine design and immunotherapy by efficiently identifying broadly reactive peptides. Our resources, including data and code, are publicly accessible at https://github.com/SkywalkerLuke/TransHLA

**Key Points:**

- We developed TransHLA, a deep learning tool for predicting epitopes across all HLA alleles using Transformer and Residue CNN architectures.
- The model uses ESM2 embeddings to improve predictive accuracy and efficiency.
- TransHLA shows superior specificity and sensitivity in identifying immunogenic epitopes and neoepitopes compared to existing models.
- Our approach offers potential advancements in vaccine design and immunotherapy through enhanced peptide analysis.

## Introduction

The intricate process of epitope presentation by human leukocyte antigen (HLA) molecules is a cornerstone of the immune system’s ability to combat pathogens, neoplasms, and its involvement in the multifaceted arenas of autoimmunity, allergies, and organ transplant rejection [1, 2]. HLA class I and II molecules play a pivotal role in presenting crucial antigen peptides to T cells[3], thereby triggering downstream immune responses.

Due to the extensive polymorphism of HLA molecules, their affinity for a wide range of peptides can vary significantly, posing a challenge for vaccine design in accurately identifying peptides that can bind to HLAs[4, 5, 6]. The burgeoning interest in HLA peptide binding has revealed the presentation of antigenic peptides by over 22,000 HLA alleles. This wealth of information constitutes a substantial database for deep learning models, offering ample resources for their development and training[7].

There are two categories of models used to predict the binding affinity between peptides and HLA alleles. The first category includes models trained individually for specific alleles, requiring users to input a peptide and select a specific allele. Examples of this category are MHCnuggets [4] and Anthem[8]. As these models assume that peptides binding to the same allele share similar characteristics, they are dependent on the alleles present in the training data, thus exhibiting limited generalization performance when applied to extensive peptides compared to the second category. The second category, known as pan-allele models, does not strictly limit predictions to specific HLA alleles. Instead, they take both the epitope and HLA allele sequences as input, as seen in models like MHCflurry [5], NetMHCpan(RRID: SCR_006604) [9], NetMHCIIpan [9], DeepSeqpanII [6], TransPHLA [10], MixMHCpred [11], and MixMHC2pred [12]. This combined embedding approach allows for a richer feature set, enhancing generalization performance. Both categories require simultaneous input of peptides and alleles to identify potential epitopes

However, while these methods can accurately determine the affinity between specific HLA residues and peptides, they rely heavily on the selection of HLA sites and lack a universal approach that focuses solely on peptides to efficiently filter out invalid peptide segments. To overcome this issue, our TransHLA does not predict binding affinity but instead predicts whether a peptide is a potential presented epitope. TransHLA only requires peptides as input and is built upon a combination of Transformer [13] and Residual CNN(Convolutional Neural Networks) architectures [14], leveraging both the sequence and structural attributes of peptides to assess the HLA binding potential. To improve feature extraction, we utilized a pretrained protein language model called Evolutionary Scale Modeling 2 (ESM2) [15], which employs an autoencoder architecture to derive a semantic and structural representation of the sequence. We have selected several state-of-the-art sequence classification models, namely TextCNN [16], TextRCNN [17], DPCNN [18], and RNN-ATTs [19], for benchmarking and comparison purposes. In addition, in the case study, we employed state-of-the-art peptide-HLA allele binding prediction software to perform predictions for all alleles. We then compared these predictions with the results obtained from TransHLA. For HLA I molecules, TransHLA achieves accuracy of 84.72% and AUC of 91.95% in IEDB test data; a 0.97% improvement in accuracy over the second-place method. For HLA II molecules, TransHLA achieves accuracy of 79.94% with an AUC of 88.14% in IEDB test data; a 2.53% improvement in accuracy over the second-place method. The comprehensive analysis of the results consistently demonstrated that TransHLA outperformed the other models in general epitope prediction for both HLA-I binding and HLA-II binding.

## Materials and methods

### Datasets

#### Data Collection

The datasets used in this work were collected and curated from IEDB(RRID: SCR_006604) [20], CEDAR [21], VDJdb [22], ImmuneCode [23], dbPepNeo2.0 [24], and NEPDB [25] databases.

The filtering criteria for each database are as follows:

**IEDB**: “MHC Ligand”, “Linear Peptide”, “Host” as Human, and “Outcome” as Positive.

**CEDAR**: “MHC Ligand”, “Linear Peptide”, “Host” as Human, and “Outcome” as Positive.

**VDJdb**: “Species” as Human.

**ImmuneCode**: We download the raw data, and add it into the analysis data.

**dbPepNeo2.0**: HC neoantigens.

**NEPDB**: “response” as positive samples.

We also removed peptides containing special characters.

#### Train-Validation-Test Data Construction

The IEDB database provided the source of our train, validation, test. The other four databases—CEDAR, VDJdb, ImmuneCode, and dbPepNeo2.0 were utilized exclusively for external test to assess the generalizability of our models. The NEPDB were utilized for assessing the performance on the neoepitope prediction. Our particular emphasis was on epitopes originating from human hosts that exhibited a positive outcome in Ligand elution/Mass spectrometry assays. For epitopes presented by HLA-II, the peptide length varied between 13 and 21 amino acids [9], whereas for epitopes presented by HLA-I, the peptide length was within the range of 8 to 14 amino acids [9].

To construct the train-validation-test datasets for both HLA-II and HLA-I, the positive samples were collected from IEDB [20] using the filtering criteria “MHC Ligand”, “Linear Peptide”, “Host” as Human, and “Outcome” as Positive. For negative samples, we used diamond [26] to blast the positive peptide sequences against the non-redundant (nr) database [27], recovering the proteins from which the sequences originated. From these proteins, random fragments excluding the positive sequences were selected to ensure non-overlap. For the positive samples, we removed peptides that did not align with any protein in the nr database using diamond.

In this way, we obtained negative samples that are representative of the potential peptide repertoire but do not include the known positive epitopes. Sequence redundancy was removed using CD-HIT [28] with a threshold of 0.8. Finally, we obtained balanced datasets for both HLA-II and HLA-I as follows: 312,245 positive samples and an equal number of negative samples for epitopes presented by HLA-II, and 459,442 positive and negative samples for epitopes presented by HLA-I. The datasets were divided into train, validation, and test sets in a ratio of 7:1:2. The details can be found in Table 1.

**Table 1.**
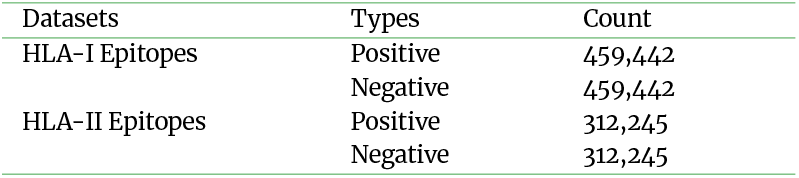
The number of samples on training datasets and independent test datasets.

To concurrently ascertain the reasonableness of our data distribution, we examined the frequencies of HLA alleles associated with peptide binding within the IEDB database. The frequency distribution of HLA alleles demonstrates highly similar characteristics across the training, validation, and test datasets (Figure 2 and Supplementary Figure 5). Our train-validation-test data includes 258 different HLA-I alleles and 227 different HLA-II alleles. Additionally, due to the larger number of HLA-A types, we plotted the top 50 alleles, while for HLA-B and HLA-C, all alleles were included. In the Supplementary Figure 5, we also plotted the top 50 alleles for HLA-DR, HLA-DQ, and HLA-DP.

**Figure 1.**
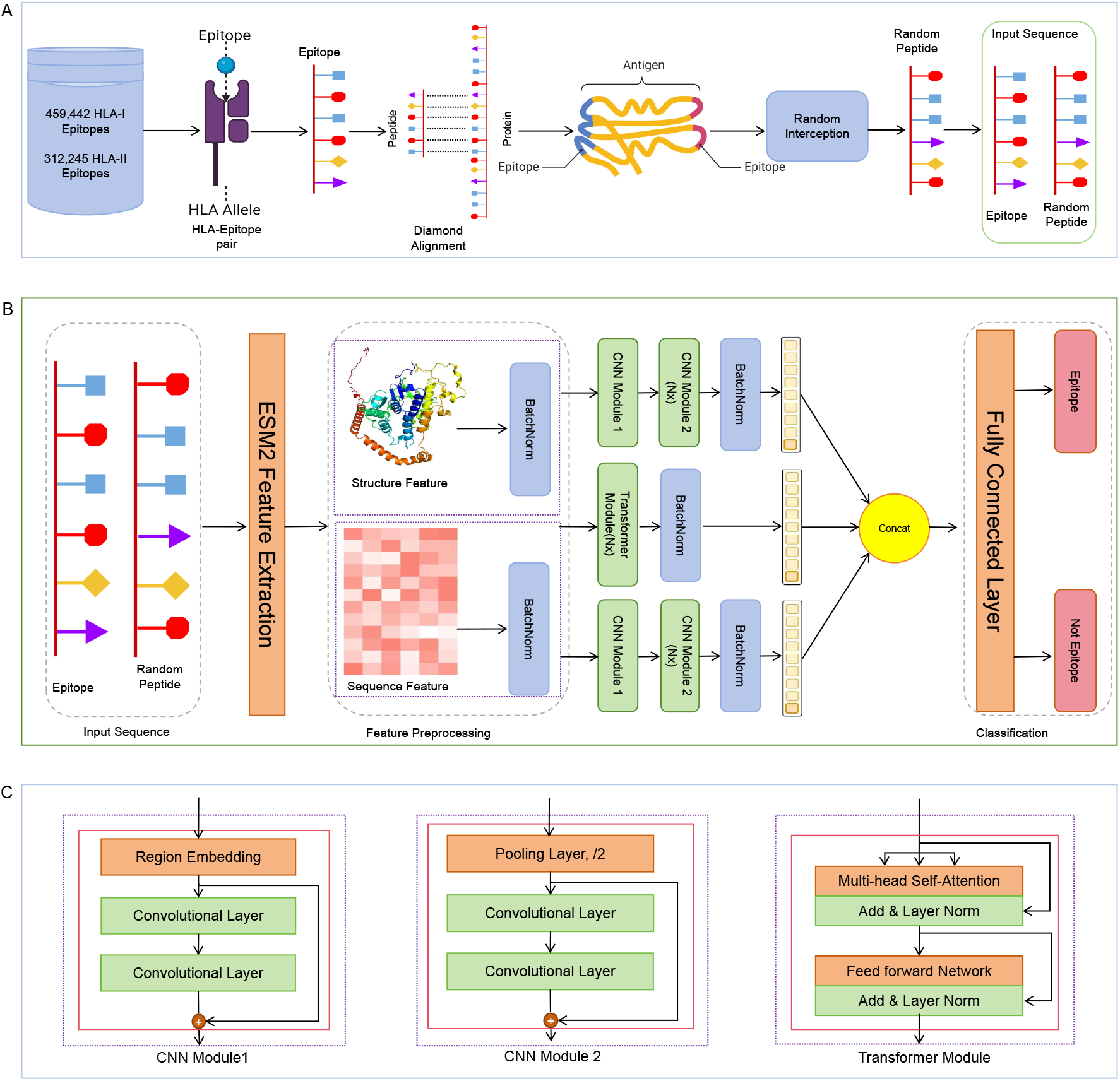
Overview of data acquisition and predictive modeling using TransHLA. (A) Data Acquisition: The dataset, derived from the IEDB, features a variety of peptide sequences that bind to HLA class I or II molecules. For negative sample generation, Non-overlapping random peptide fragments were sourced by matching positive peptides to their originating proteins through sequence alignment. Then, the Non-overlapping random peptide fragments were processed with CD-HIT to achieve a reduced redundancy, resulting in the final set of negative samples. (B)With ESM2’s advanced modeling capabilities, we generated high-dimensional sequence embeddings for the peptides associated with both HLA classes. Concurrently, structural insights were obtained through ESM2’s contact map predictions, yielding structure embeddings. These two distinct yet complementary sets of embeddings were crafted to capture the intricate nature of peptide-HLA interactions. describes the process from data input to epitope presentation prediction. (C) The architecture of the different modules, including the CNN Module 1, CNN Module 2 and the Transformer module.

**Figure 2.**
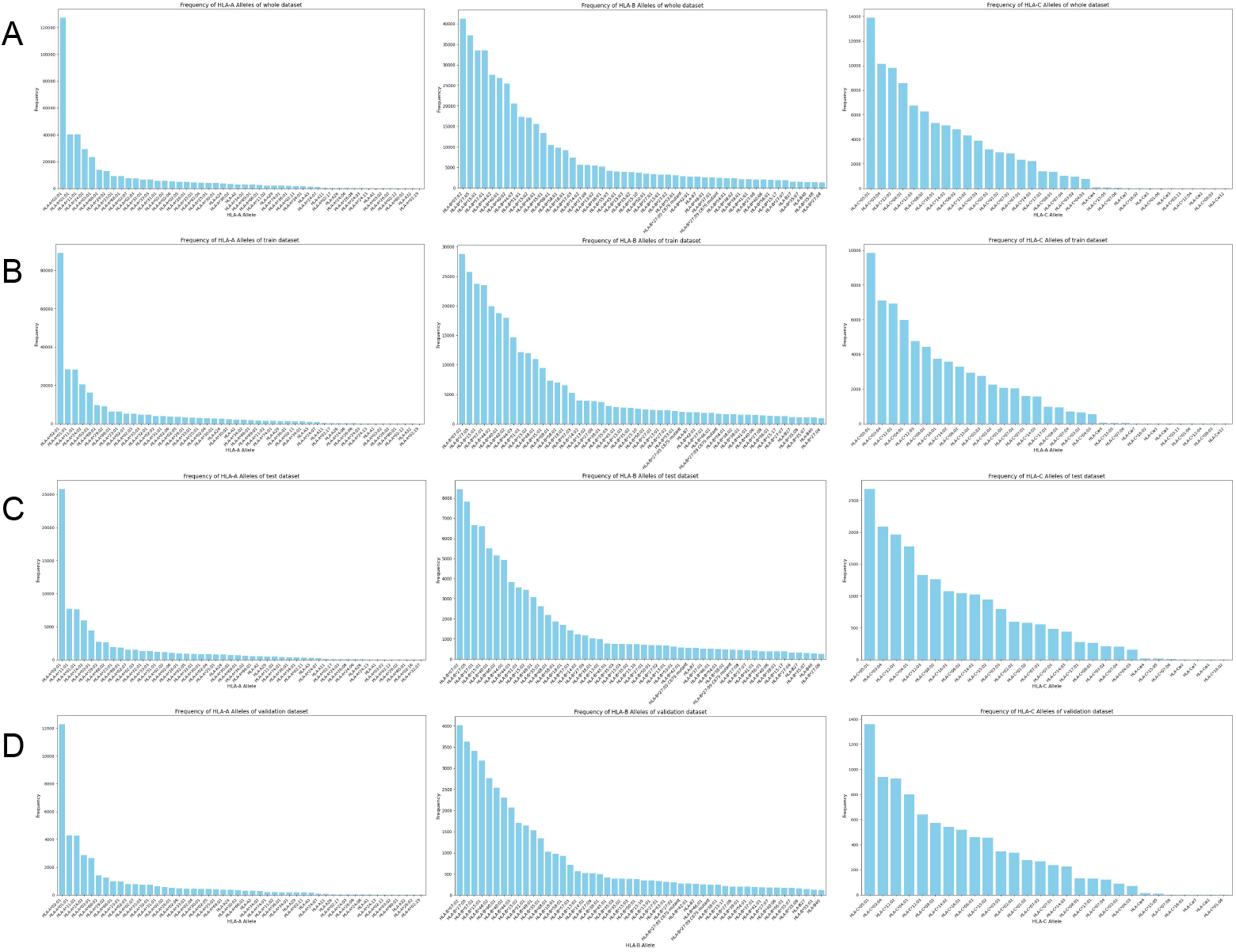
This figure illustrates the experimental distribution of major HLA-I alleles in the IEDB. We plotted the alleles by type, where subfigure A shows the distribution for all data, B for the training dataset, C for the test dataset, and D for the validation dataset. Left represents HLA-A alleles, middle represents HLA-B alleles, and right represents HLA-C alleles.

### Pre-trained Embeddings for Sequence and Structure

Pre-trained protein language models[29, 15, 30] have been extensively applied in various tasks, such as protein classification, by providing intricate representations of protein sequences [31, 32, 33]. Additionally, AlphaFold2 [34] and ColabFold [35] have set high standards in protein structure prediction.

In our approach, we address the issue of HLA-I binding epitopes with lengths less than 14 and HLA-II binding epitopes with lengths less than 21 by padding the end with ones. This padding technique ensures that the sequences have the required lengths. Subsequently, we utilize the ESM2 protein language model to extract sequence embeddings and predict structure embeddings for these epitopes[15].

### The Architectures of the Deep Learning Model

#### The Transformer module

To enhance the extraction of global features, we incorporated the Transformer encoder module [13, 36], which utilizes inputs in the form of pre-trained sequence features extracted by ESM2, represented as a **E** ∈ *R*^1280*×peptide*_*length*^ matrix. The module leverages a multi-head attention mechanism to facilitate effective global feature extraction.

Within each attention head, three key components are involved: *Q* (query), *K* (key), and *V* (value). *Q* represents the current position being attended, while the *K* and *V* represent other positions in the peptide sequence. By computing the attention weights between the *Q* and *K*, the model determines the importance of each position and assigns higher weights to more relevant positions. The values are then combined based on these attention weights to generate the output representation, and the scaled dot-product attention is calculated as:

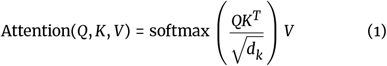

*d*_*k*_ is the dimension of the key vectors.

The multi-head attention is achieved through a series of operations to transform the input vectors *Q, K*, and *V* for *h* times (where *h* is the number of heads). Each transformed vector undergoes scaled dot-product attention independently. Finally, the attention outputs are concatenated and further transformed. This process can be expressed as follows:

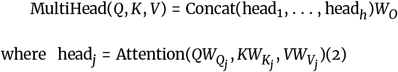

In the given equation, 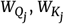, and 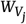 represent weight matrices for each head corresponding to the *Q, K*, and *V* vectors, respectively. *W*_*O*_ denotes the weight matrix for the output. This formulation allows for weight sharing across heads, reducing redundancy and promoting a more compact representation.

#### The CNN module

To enhance feature extractions, TransHLAs employs two structurally identical CNNs, each consisting of a CNN module 1 for region embedding, followed by multiple layers of CNN module 2. These modules process both the pre-trained sequence features **E** and the contact map structural features extracted by ESM2, with the contact map being a symmetric matrix **S** ∈ *R*^*peptide*_*length×peptide*_*length*^. Residual connections are implemented between each module to prevent gradient vanishing and ensure effective training of the deep network structure.

##### CNN Module 1

It first applies a text region embedding layer to get a dense representation of sequences:

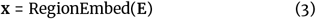

where **x** ∈ *R*^*M×D*^ and *M* is the number of text regions and *D* is the region embedding dimension.

This is followed by a convolution block, which contains two convolutional layers each with 256 feature maps:

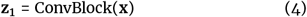

where **z**_1_ ∈ *R*^*M×*256^.

##### CNN Module 2

Following the CNN module 1, the CNN module 2 commences with a pooling layer that reduces the length of the feature map to half of its original size. Subsequently, CNN module 2 employs a convolutional block with the same structure as the one in CNN module 1, featuring an isometric convolutional layer with 256 channels.

**z**_1_:

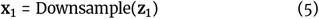

where **z**_2_ ∈ *R*^*M*/2*×*256^.

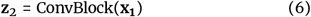

This is repeated for *L* times, with the downsampling layer and *l*-th CNN Modules being:

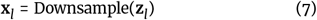

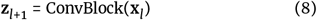

where 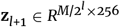.

After the final Downsample layer, we add a batchnorm layer, which helps reduce internal covariate shift and acts as a regularization technique.

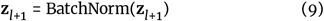

#### The TIM Loss

The TransHLA framework is developed based on a modified Transductive Information Maximization (TIM) loss function, as introduced by Boudiaf et al. (2020) [37], which merges conventional cross-entropy with a mutual information component,tailored for empirical analysis. We address the empirical mutual information within dataset *X* (comprising amino acid sequences) linked to their respective outcomes *Y* (which now signify epitope presentation). The first factor is the empirical conditional entropy of the outcomes given the data, denoted as *H*(*Y*|*X*), which reduces uncertainty in predictions for unlabeled samples, encouraging the model to output confident predictions. The next factor is the empirical marginal entropy of the outcomes, denoted as *H*(*Y*), which encourages a uniform distribution of labels, preventing bias towards any particular class. To further calibrate the binary classification process, the cross-entropy loss, indicated as CE, between the model’s predictions and the actual outcomes is incorporated. By integrating these entropy terms, the TIM loss enhances the model’s generalization in few-shot scenarios without the need for complex meta-learning schemes. The formulation for these components is defined as follows

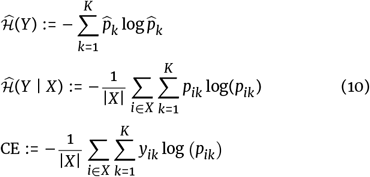

Let |*X*| denote the total count of sequences within the dataset, with *i* being the sequence identifier in *X*, and *K* representing the possible outcome categories. The variable *p*_*ik*_ is the predicted like-lihood of the *i*-th amino acid sequence being classified within the *k*-th category. The binary variable *y*_*ik*_ is used to indicate if the *i*-th sequence is actually categorized under class *k*. We assign *K* = 2 for this model since the task at hand is a binary classification problem.

The final loss function for TransHLA is defined as:

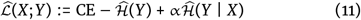

Where α is hyperparameter that determine the rate of convergence for each term in the loss function. In experiments, we set α = 0.04, considering the standard cross-entropy loss and standard mutual information.

By selecting these particular hyperparameter values, we maintain fairness and impartiality in TransHLA’s training process.

### Performance evaluation

In the Benchmark Results with other sequence classification models, we employed the aforementioned metrics, including accuracy (ACC), Recall, F1-score (F1), and Matthews Correlation Coefficient (MCC), to evaluate the performance of TransHLA.

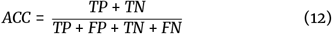

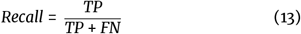

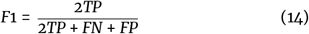

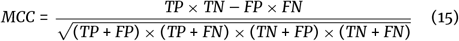

Additionally, in the Comparison Results with other HLA-epitope binding software across extensive datasets, we augmented our assessment with two further metrics, Precision and Specificity. These additional indicators highlight how our methodology has overcome the limitations commonly associated with traditional affinity-binding software, particularly the tendency to incorrectly classify negative samples as positive.

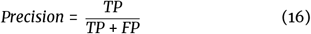

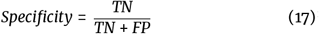

Where TP (true positives) represents the number of correctly identified true epitope, TN (true negatives) represents the number of correctly identified the normal peptide, The FP (false positives) represents the number of instances where normal peptides were incorrectly identified as epitopes, and FN (false negatives) represents the number of instances where epitopes were incorrectly identified as normal peptides.

In addition to these metrics, we also utilize Receiver Operating Characteristic (ROC) and Precision-Recall (PR) curves as significant evaluation tools for classification accuracy. The Area Under the ROC Curve (AU-ROC) and the Area Under the Precision-Recall Curve (AU-PRC) values quantify the overall performance by measuring the area beneath the ROC and PR curves, respectively.

## Results

### Benchmark Results with other sequence classification models

Since this paper focuses on the epitope presentation classification problem, a corresponding software for comparison is not yet established. We conducted comparison experiments on independent test sequences from IEDB including 92,347 HLA-I binding epitopes, 65,105 HLA-II binding epitopes and 157,879 random sequences with the state-of-the-art sequence classification models, including TextCNN [16], TextRCNN [17], DPCNN [18], and RNN-ATTs [19].

The performance metrics are presented in Table 2. TransHLA exhibits enhanced performance in classifying HLA-I epitopes across five metrics, including ACC, F1, Recall, MCC and AUC(AU-ROC). The corresponding values for TransHLA are 0.847, 0.846, 0.694, and 0.920, respectively. In comparison, the second-ranked software obtains scores of 0.838, 0.838, 0.68 and 0.910 for the same metrics, respectively. Figure 3A and 3C illustrate the ROC and PR curves of the compared models.

**Table 2.** Benchmark Results with other sequence classification models.

| Type | Method | ACC (%) | F1(%) | Recall(%) | MCC | AUC (%) |
| --- | --- | --- | --- | --- | --- | --- |
| HLA-I | TransHLA | **84.72** | **84.59** | 83.92 | **0.69** | **91.95** |
|  | TextCNN | 81.63 | 79.61 | 83.50 | 0.63 | 89.37 |
|  | TextRCNN | 81.21 | 76.65 | 83.78 | 0.64 | 87.62 |
|  | DPCNN | 83.75 | 83.76 | **83.96** | 0.68 | 90.97 |
|  | RNN-ATTs | 81.17 | 82.27 | 81.46 | 0.63 | 87.98 |
| HLA-II | TransHLA | **79.94** | **81.07** | **86.19** | **0.60** | **88.14** |
|  | TextCNN | 73.26 | 73.49 | 72.64 | 0.47 | 80.64 |
|  | TextRCNN | 70.96 | 72.02 | 69.28 | 0.42 | 78.21 |
|  | DPCNN | 77.41 | 77.91 | 75.98 | 0.55 | 85.30 |
|  | RNN-ATTs | 69.04 | 69.83 | 67.89 | 0.38 | 75.81 |

**Figure 3.**
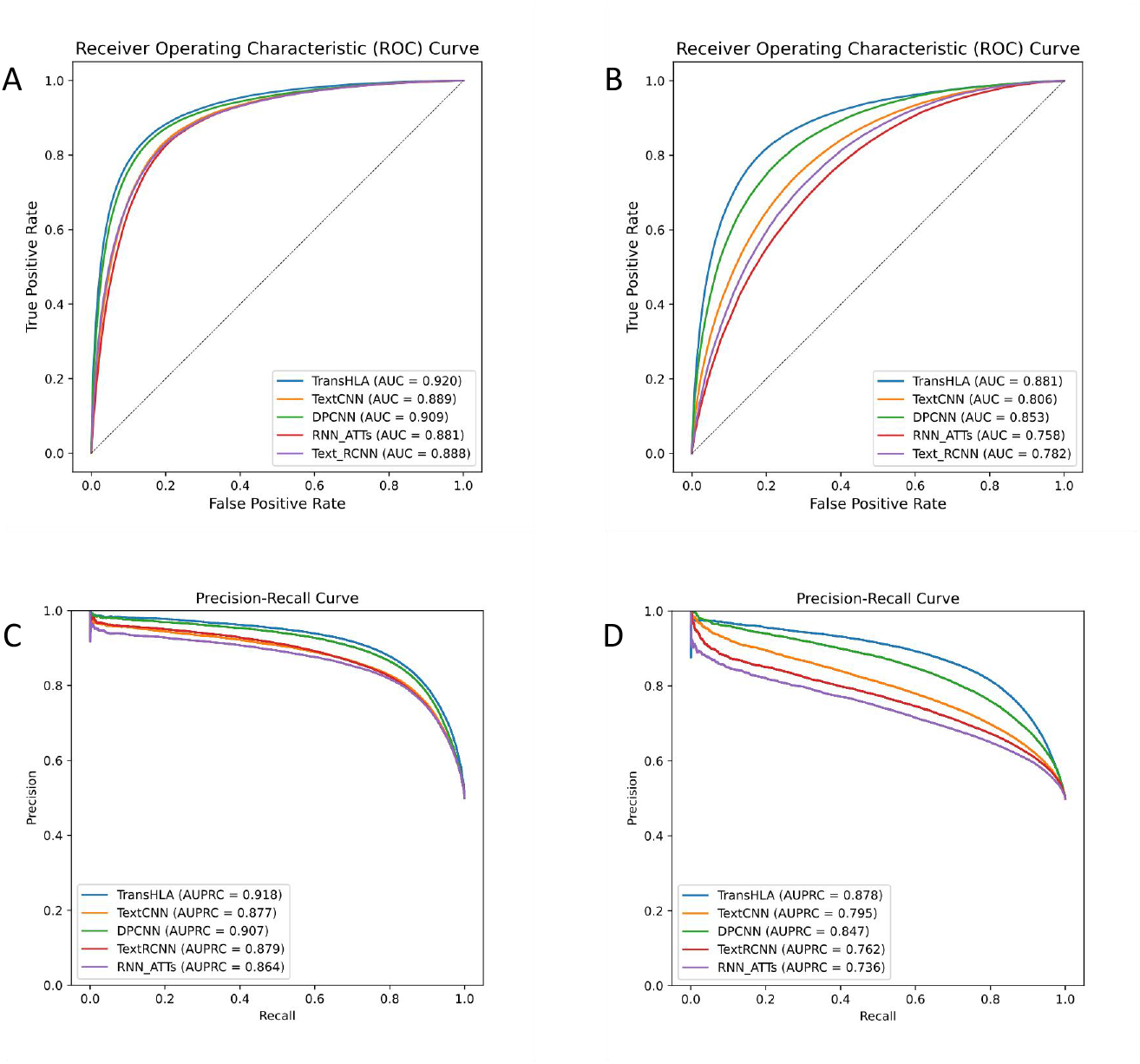
This figure evaluates TransHLA’s epitope prediction capabilities, benchmarked against other models using two key statistical metrics: AUROC and AUPRC. Subfigures (A) and (C) examine HLA-I class performance, with (A) showing AUROC and (C) presenting AUPRC. The graphs demonstrate TransHLA’s proficiency in distinguishing between epitope and non-epitope peptides for HLA-I, where AUROC indicates its discriminative power and AUPRC reflects the precision-recall trade-off. Subfigures (B) and (D) extend the analysis to HLA-II, with (B) displaying AUROC and (D) illustrating AUPRC. The performance depicted emphasizes TransHLA’s effectiveness in identifying HLA-II epitopes, highlighting its ability to differentiate between classes under imbalanced distributions. Together, the subfigures demonstrate TransHLA’s advantage over conventional models in epitope prediction for both HLA classes.

HLA-II binding predictions are known to be more complex compared to HLA-I predictions. [38, 39, 40, 12] Consequently, the performance of the models, in general, is inferior in terms of HLA-II binding metrics of other models, with an average decrease of 9.3% in ACC, 7.26% in F1, 11.73% in Recall, 0.19 in MCC, 9.00% in AUC. However, even in the challenging task of HLA-II binding prediction, TransHLA demonstrates robust classification performance. Compared to the values achieved in HLA-I binding prediction, TransHLA only experiences a decrease of 4.78%in ACC, 3.52% in F1, 0.09 in MCC, and 3.81% in AUC. Remarkably, TransHLA achieves an increase of 2.27% in Recall. Furthermore, in the prediction of HLA-II epitope binding, TransHLA demonstrates superior performance across all evaluation metrics. Compared to the next best-performing models, TransHLA achieves an improvement of 0.25 in ACC, an enhancement of 0.316 in the F1 Score, a boost of 1.02 in Recall, and a substantial improvement of 0.548 in MCC.

### The prediction performance in different alleles

To further discuss TransHLA’s performance across different alleles, we calculated the positive sample prediction accuracy, or recall, for alleles sorted by the number of corresponding epitopes from high to low. The detailed results are shown in Supplementary Figure 6. Overall, when frequencies are higher, prediction performance tends to improve. For example, in HLA class I, when the frequency is above 42, the average performance is around 80%, some achieves 90%,such as HLA-B*40:01, HLA-B*44:03. However, when the frequency is below 42, some alleles show fluctuating results, with accuracy for certain alleles dropping to below 55%, such as HLA-B41:05, HLA-B41:04, and HLA-B15, leading to lower overall effectiveness compared to class II. In HLA class II, when frequencies exceed 200, the overall accuracy is higher, about 80% or more. For frequencies between 100 and 200, such as HLA-DRA01:01/DRB107:01 to HLA-DQA102:01/DQB03:03, performance is slightly lower, with accuracy around 50-65%. However, even alleles with fewer epitopes can still achieve relatively high accuracy. In class I, examples include HLA-B41:05, HLA-B41:04, and HLA-B15, while in class II, examples are HLA-DRB*03:01 and some alleles with even lower frequencies.

Meanwhile, due to the random allocation of epitopes to train-validation-test sets, some test epitopes’ alleles did not appear in the train and validation sets. These data were also included in our analysis. For class I, epitopes such as ‘ALWGFFPVL’, ‘LDTNADKQLSF’, ‘WQQGLRVSF’, ‘ILDTAGKEEY’ were included. For class II, epitopes like ‘PKYVKQNTLKLAT’ and ‘ISTNIRQAGVQYSRA’ were analyzed. We found that our model predicted these peptides as positive samples with probabilities greater than 0.8. The specific prediction results are in Supplementary Table 3, and the corresponding alleles are listed in Supplementary Table 4.

### Comparison Results with other HLA-epitope binding software in the case study

We employed our software along with various state-of-the-art epitope-HLA binding prediction tools, including Mhcflurry [5], NetMHCpan4.1b [9], NetMHCIIpan4.3b [9], TransPHLA [10], Anthem[8], MixMHCpred [11], DeepSeqPanII [6], MixMHCIIpred [12] and Mhcnuggets [4] to evaluate their accuracy in correctly identifying sequence as potential epitopes from CEDAR [21], VD-Jdb [22], ImmuneCode [23], and dbPepNeo2.0 [24] datasets. Our analysis yielded a total of 21,387 HLA-I binding epitopes and 3,580 HLA-II binding epitopes in the case study. Our criterion for deeming a peptide as a presentation-worthy epitope is that it must exhibit binding affinity to at least one major HLA allele. Mhcflurry contains 11,576 HLA-I alleles, Mhcnuggets contains 118 HLA-II alleles and 106 HLA-I alleles, and DeepSeqPanII contains 61 HLA-II alleles. And the details information of alleles used in each tool can be found in our Data availability.

In this case study, we added random sequences in the same quantity as the identified persistent epitopes. The results of the experiments are presented in Table 3. For the prediction of HLA-I binding, we compared the performance of the TransHLA against Mhcflurry, NetMHCpan, MixMHCpred, TransPHLA, Anthem, and Mhcnuggets. In the prediction of HLA-II binding, we compared the performance of the TransHLA, DeepSeqPanII, Mhcnuggets, MixMHC2pred, NetMHCIIpan, and MixMHC2pred. The details of the parameters used in the mentioned methods can be found in the Supplementary File. TransHLA, Mhcflurry, and NetMHC-pan demonstrated good performance in predicting HLA-I binding. Among the seven models, TransHLA achieved the highest ACC of 83.09%, precision of 85.22%, and specificity of 86.11%, followed by Mhcflurry with 82.96%, 81.95%, and 81.38% in the same metrics. Besides, TransHLA, in conjunction with the Mhcflurry and NetMHCpan, obtained the highest MCC of 0.66 in this experiment. Mhcflurry had the highest F1-score of 83.24%, while TransHLA obtained the second highest F1-score of 82.96%. Upon closer examination of the F1 score metrics, it was observed that TransHLA attained the highest true negative (TN) rate of 86.1%, which surpassed that of MhcFlurry and NetMHCpan at 81.4% and 76.2%. Conversely, NetMHCpan exhibited the highest true positive (TP) rate at 86.0%. This differential performance indicates that TransHLA demonstrates a more robust capability in filtering out noise during epitope identification. In the prediction of HLA-II binding, Tran-sHLA showed better performance than other models, achieving the highest scores in ACC (66.88%), F1 (64.08%), MCC (0.34), Precision (66.99%), and Specificity (74.67%). In contrast, while Mhcnuggets achieved a high recall of 89.69% in HLA-II binding, its precision was only 50.04%, indicating that it misjudged a large number of negative samples as epitopes. This causes a lot of noise to be mixed into the screening results. NetMHCIIpan 4.3b and MixMHC2pred achieved ACC scores of 60.34% and 61.62%, respectively on HLA-II, but they have much lower Specificity scores than TransHLA. DeepSeqPanII performs worse than the other models in HLA-II binding with 49.68% ACC. To further validate TransHLA’s robust-ness in real-world scenarios with more negative than positive samples, we conducted an experiment with a 1:4 positive-to-negative sample ratio. TransHLA achieved an ACC of 84.75% and Specificity of 86.11% for HLA-I binding, and an ACC of 71.14% and Specificity of 74.12% for HLA-II. Detailed results are in Supplementary Table 2.

### TransHLA extracts a high-quality peptide embedding in low-dimension

To assess the feature extraction capability of the model, we under-took dimension reduction and visualization of the penultimate layer features derived from the self-trained models outlined in test data, along with random embeddings and the TransHLA embeddings. The PCA [41] layouts of the learned representations for HLA-I epi-topes binding prediction (Figure 4).

**Figure 4.**
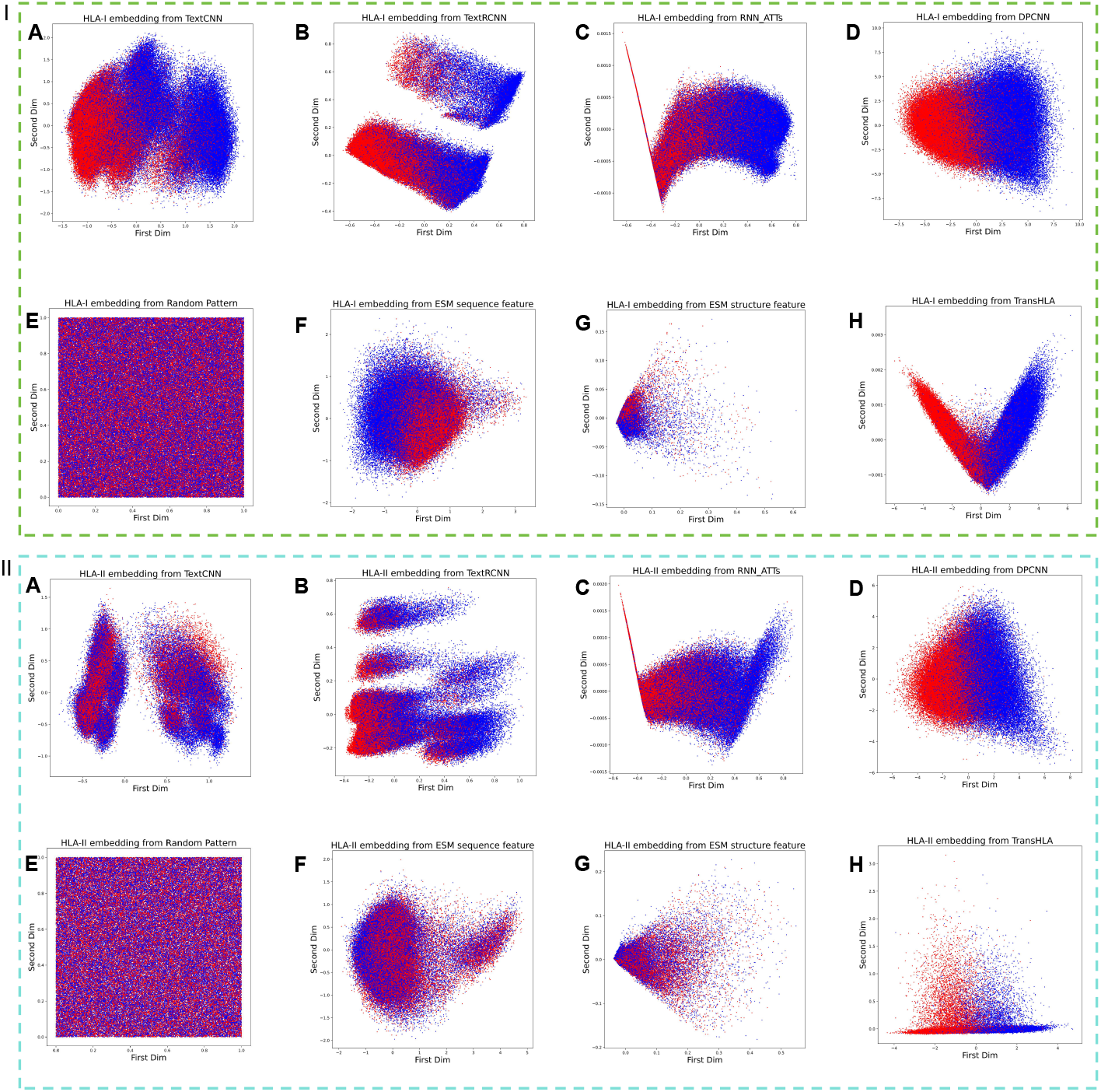
This figure displays the PCA-based 2D feature space distribution of HLA-I and HLA-II epitope presentation test sets. The space is split into two regions: Region I for HLA-I epitopes and Region II for HLA-II epitopes. Each region shows positive samples as blue dots and negative samples as red dots, with representations from various model embeddings (A-H) including Random Pattern, TextCNN, TextRCNN, RNN-ATTs, DPCNN, random initial, ESM2 pre-trained sequence, structure pre-trained, and TransHLA.

**Figure 5.**
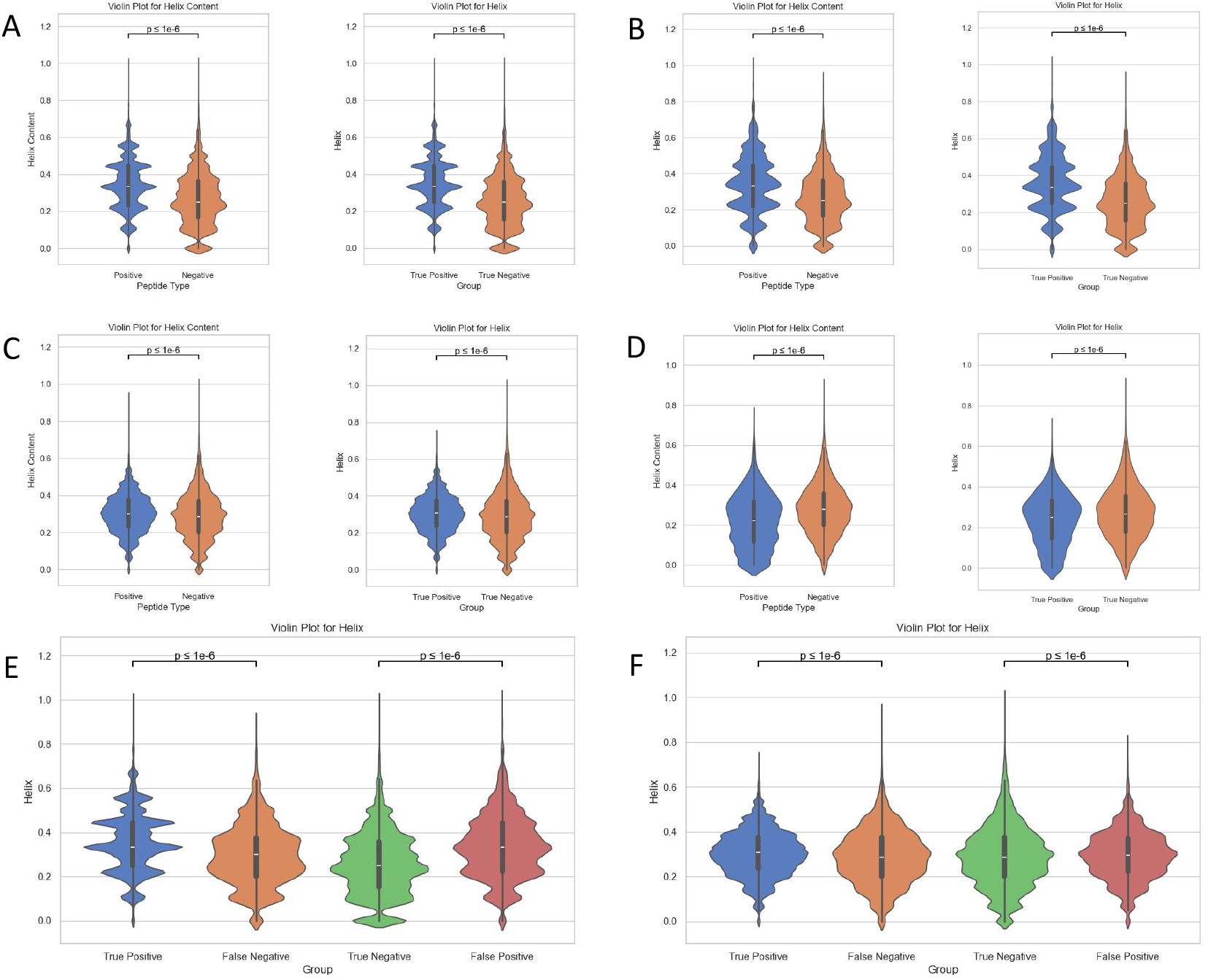
This comprehensive figure presents a series of violin plots illustrating the ‘helix content’ structure property of peptides across various sample subsets for HLA-I and HLA-II molecules. Subfigure (A) delineates the helix content distribution in independent test samples for HLA-I, separated into positive and negative samples, with each subgroup’s statistical significance assessed via t-tests and annotated with corresponding p-values. Subfigure (C) mirrors this setup for HLA-II independent test samples, highlighting the comparative helix content distributions. The external dataset distributions for HLA-I and HLA-II are respectively showcased in subfigures (B) and (D), emphasizing the metric’s external validity. Subfigures (E) and (F) delve deeper, contrasting the helix content of true positives and false negatives against true negatives and false positives within HLA-I and HLA-II datasets, respectively.

When comparing with random embedding, both DPCNN (Figure 4 I-D, II-D) and TransHLA (Figure 4 I-H, II-H)demonstrate superior embedding effects on HLA-I and HLA-II binding epitopes, thereby exhibiting distinct discrimination between positive and negative samples. However, in the low-dimensional visualization of DPCNN, a considerable number of positive and negative samples overlap at the junction, while TransHLA exhibits a more apparent boundary in comparison.

### Good performance achieved in TCR experiments validated NeoEpitope prediction by TransHLA

To validate the effectiveness of TransHLA in NeoEpitope prediction, we employed newly identified epitopes with immunogenicity verified by TCR experiments from NEPDB [25] as positive samples and compared our software with several other widely used tools for NeoEpitope prediction, including NetMHCpan, Mhcflurry, Mhc-nuggets, MixMHCpred, TransPHLA, and Anthem. In NEPDB, we collected a total of 139 neoantigens presented by HLA-I alleles. Additionally, we used a 1:1 ratio of randomly selected negative samples to form our final dataset.

The results of the comparison between these tools are presented in Table 4. Among the compared methods, TransHLA and Mhcflurry achieved the top two performances. Specifically, TransHLA attained the highest accuracy of 90.65%, followed closely by Mhcflurry, which reached an accuracy of 89.56%. Furthermore, although NetMHCpan4.1b achieved a recall of 98.56%, its specificity was only 71.94%, highlighting its tendency to misclassify negative samples as positive. In contrast, TransHLA maintained a strong balance between recall and specificity, achieving a specificity of 87.77% and a high recall of 93.52%. Interestingly, all models demonstrated higher recall in prediction compared to MHC experimental validation, with epitopes validated by TCR experiments (Table 4) performing significantly better in predictions than those validated solely by MHC experiments (Table 3). TransHLA demonstrated the highest Specificity, Precision, ACC, F1, and MCC, ensuring that it does not miss a large number of immunogenic epitopes while effectively filtering out the majority of negative samples.

**Table 3.**
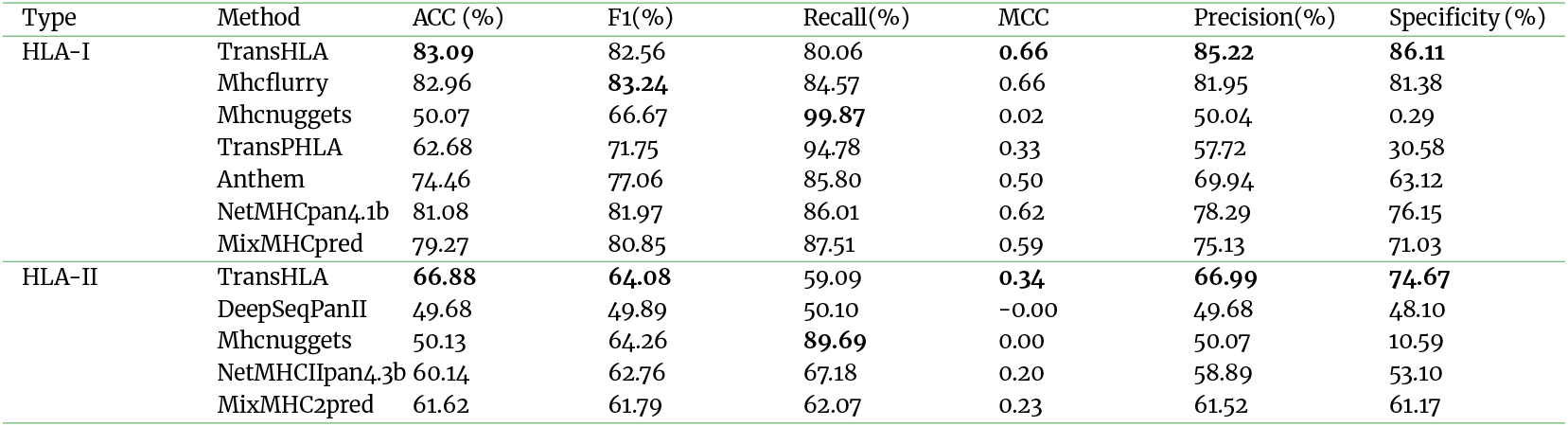
Comparison Results with other HLA-epitope binding softwares in External Epitope Datasets Case Study.

| Type | Method | ACC (%) | F1(%) | Recall(%) | MCC | Precision(%) | Specificity (%) |
| --- | --- | --- | --- | --- | --- | --- | --- |
| HLA-I | TransHLA | **83.09** | 82.56 | 80.06 | **0.66** | **85.22** | **86.11** |
|  | Mhcflurry | 82.96 | **83.24** | 84.57 | 0.66 | 81.95 | 81.38 |
|  | Mhc nuggets | 50.07 | 66.67 | **99.87** | 0.02 | 50.04 | 0.29 |
|  | TransPHLA | 62.68 | 71.75 | 94.78 | 0.33 | 57.72 | 30.58 |
|  | Anthem | 74.46 | 77.06 | 85.80 | 0.50 | 69.94 | 63.12 |
|  | NetMHCpan4.1b | 81.08 | 81.97 | 86.01 | 0.62 | 78.29 | 76.15 |
|  | MixMHCpred | 79.27 | 80.85 | 87.51 | 0.59 | 75.13 | 71.03 |
| HLA-II | TransHLA | **66.88** | **64.08** | 59.09 | **0.34** | **66.99** | **74.67** |
|  | DeepSeqPanII | 49.68 | 49.89 | 50.10 | -0.00 | 49.68 | 48.10 |
|  | Mhc nuggets | 50.13 | 64.26 | **89.69** | 0.00 | 50.07 | 10.59 |
|  | NetMHCIIpan4.3b | 60.14 | 62.76 | 67.18 | 0.20 | 58.89 | 53.10 |
|  | MixMHC2pred | 61.62 | 61.79 | 62.07 | 0.23 | 61.52 | 61.17 |

**Table 4.** Comparison Results with other HLA-epitope binding software in NeoEpitope Prediction Case Study.

| Type | Method | ACC (%) | F1(%) | Recall(%) | MCC | Precision(%) | Specificity (%) |
| --- | --- | --- | --- | --- | --- | --- | --- |
| HLA-I | TransHLA | **90.65** | **90.91** | 93.52 | **0.81** | **88.43** | **87.77** |
|  | MHCflurry | 89.56 | 90.23 | 96.40 | 0.80 | 84.81 | 82.73 |
|  | Mhc nuggets | 50.00 | 66.67 | **100.00** | 0.00 | 50.00 | 0.00 |
|  | TransPHLA | 64.03 | 73.40 | 99.28 | 0.40 | 58.23 | 28.78 |
|  | Anthem | 82.01 | 84.08 | 94.96 | 0.66 | 75.43 | 69.06 |
|  | NetMHCpan4.1b | 85.25 | 86.98 | 98.56 | 0.73 | 77.84 | 71.94 |
|  | MixMHCpred | 84.89 | 86.36 | 95.68 | 0.71 | 78.70 | 74.10 |

### Ablation study

To evaluate the contribution of different components of TransHLA to its performance on test data, we conducted an ablation study on seven variants: without the transformer module, omitting structure pre-trained embedding, removing sequence pre-trained embedding, deleting the CNN module, without any pre-trained embedding, changing TIM loss to Cross Entropy loss, and randomly extending the peptide sequence by 0-3 units on each end using proteins aligned with Diamond. The performance metrics for different modules across both HLA-I and HLA-II models are illustrated in Table 5.

**Table 5.** Ablation Study on different modules.

| Type | Method | ACC (%) | F1(%) | Recall | MCC | AUC (%) |
| --- | --- | --- | --- | --- | --- | --- |
| HLA-I | TransHLA | **84.72** | 84.59 | 83.92 | **0.69** | **91.95** |
|  | w/ Transformer module | 84.18 | 83.89 | 82.39 | 0.68 | 91.26 |
|  | w/ CNN module | 83.43 | 83.09 | 84.85 | 0.67 | 90.70 |
|  | w/ structure pre-trained embedding | 83.78 | **84.80** | 81.85 | 0.68 | 91.39 |
|  | w/ sequence pre-trained embedding | 73.37 | 73.45 | 73.56 | 0.4782 | 80.72 |
|  | w/ any embedding | 84.56 | 84.68 | **84.87** | 0.68 | 91.33 |
|  | w/ TIM Loss | 84.04 | 83.86 | 82.98 | 0.68 | 90.93 |
|  | extend sequence | 75.60 | 75.98 | 77.21 | 0.51 | 83.20 |
| HLA-II | TransHLA | **79.94** | **81.07** | **86.19** | **0.60** | **88.14** |
|  | w/ Transformer module | 79.38 | 79.42 | 79.31 | 0.59 | 87.31 |
|  | w/ CNN module | 74.43 | 74.09 | 74.85 | 0.49 | 82.36 |
|  | w/ structure pre-trained embedding | 79.68 | 80.47 | 79.41 | 0.59 | 87.49 |
|  | w/ sequence pre-trained embedding | 67.52 | 61.21 | 51.41 | 0.37 | 76.84 |
|  | w/ any embedding | 78.35 | 78.44 | 77.87 | 0.57 | 86.13 |
|  | w/ TIM Loss | 79.14 | 78.14 | 74.79 | 0.58 | 86.56 |
|  | extend sequence | 77.63 | 77.34 | 78.07 | 0.55 | 85.02 |

Based on the comparisons conducted, it is evident that the sequence embedding approach is significantly more effective in capturing the peptide features, achieving 84.56% and 79.68% accuracy in HLA-I and HLA-II, respectively. However, the extended sequences only achieved 75.60% and 77.63% accuracy in HLA-I and HLA-II, respectively. This may be due to the fact that, after epitope extension, the distinguishing features are no longer as prominent. Epitopes are a specific type of peptide with unique overall chemical and structural properties, such as hydrophobicity, aromaticity, and secondary structure. Although the extended sequences include the original fragments, they can be considered as new peptide fragments that may no longer retain the original characteristics of the epitope, thus potentially confusing the model [42, 43, 44].

The CNN modules, due to their difference in global feature extraction compared to pretrained transformers, contribute substantially to performance enhancement, achieving 84.18% and 79.38% accuracy in HLA-I and HLA-II. Additionally, other modules also play their respective roles in the overall efficacy of the system. This is corroborated by ablation studies, which demonstrate that each module contributes positively to the model’s predictive capabilities.

### Chemical and Secondary Structure Patterns of Epitopes in Antigen Presentation

We further utilized Biopython [45] to investigate the chemical properties and secondary structure of the peptide segments. The analyzed chemical properties included aromaticity, hydrophobicity, flexibility, instability index, isoelectric point, average molecular weight, and mean charge at pH 7. For flexibility, we used a window size of 9-mer length [46] with a sliding step of 1, and averaged the resulting values. For peptides with a length less than 9, a flexibility value of 0 was assigned. Secondary structure analysis covered helix, sheet, and coil content. Using XGBoost [47] for feature selection on the same dataset as TransHLA, we identified the top three features for HLA-I as helix content, aromaticity, and flexibility, while for HLA-II, the top three were aromaticity, helix content, and hydrophobicity (Supplementary Table 1). We further analyzed these features in epitopic vs. non-epitopic regions and compared feature distributions between true positives and true negatives predicted by TransHLA.

Through rigorous statistical analyses conducted on our test dataset we made some intriguing observations regarding the helix content of epitopes and non-epitopes. For helix content feature, our findings demonstrate that epitopes consistently exhibit higher helix content in comparison to non-epitopes (Figure 5 A left, C left) with a statistically significant p-value of less than 1 *×* 10^−6^. Furthermore, we also examined the helix content of true positive and true negative samples predicted by TransHLA and found a similar trend (Figure 5 A, C) with a p-value of less than 1 *×* 10^−6^ (Figure 5 Aright, C right). Aromaticity exhibits a similar pattern to helix content, where epitopes are significantly higher than non-epitopes(Supplementary Figures 4 A, C). Upon conducting a comparative analysis of Figure 5 B and D, a notable divergence was observed when testing the model on the case study dataset. Specifically, epitopes associated with HLA Class I consistently exhibited their characteristic higher helix content, as evidenced by a statistically significant p-value of less than 1 *×* 10^−6^. Conversely, in the case of epitopes associated with HLA Class II, this characteristic was not maintained, as indicated that the helix content of the epitopes is lower than non-epitopes. Consequently, the performance of TransHLA declined compared to the benchmark reported in test data. Moreover, from Supplementary Figure 4 B and D, we can see that aromaticity, which is related to structural stability[48, 49], following a similar trend with helix content Feature. In Supplementary Figures 3C and D, HLA II epitopes show lower hydrophobicity compared to HLA I epitopes and non-epitopes, aligning with their role in presenting exogenous antigens in aqueous environments. This hydrophilicity facilitates hydrogen bonding with water, stabilizing peptide structures [50, 51, 52]. As for flexibility, due to the special handling of this feature, we only calculated this feature for peptides 10 residues or longer, and found it to have limited statistical utility (Supplementary Figure 2). Flexibility showed higher importance in HLA-I epitopes due to shorter peptides (8-9 residues) being assigned a value of zero, artificially inflating its relevance.

Further exploring the positive epitope samples, which include True Positives (TP) and False Negatives (FN), as well as the negative epitope samples, which include True Negatives (TN) and False Positives (FP), we made an intriguing observation (Figure 5 E, F). There is a significant distribution gap in helix content between the correctly classified samples (TP and TN) and the incorrectly classified samples (FP and FN) predicted by TransHLA (*p* - *value* ≤ 1 *×* 10^−6^).

Interestingly, the incorrectly classified samples (FN and FP) exhibit relatively similar flexibility distributions. This similarity in epitope characteristics between the FN and FP samples likely contributes to the high difficulty in accurate prediction. Similarly, from Supplementary Figures 3, and 4 (E, F), we can observe that Hydrophobicity, and Aromaticity, exhibit similar conclusions.

Statistical data indicates that both HLA-I and HLA-II epitopes are more rigid and stable, supporting the notion that rigidity facilitates epitope recognition [48, 49, 53, 54]. Our model shows that peptide segments with higher helix content and aromaticity are more likely to be recognized as epitopes in both classes. A key distinction lies in hydrophobicity—HLA-I epitopes are more hydrophobic, while HLA-II epitopes are more hydrophilic, likely reflecting their roles in presenting intracellular and extracellular antigens, respectively, with the latter often exposed to aqueous environments.

## Conclusion and Discussion

In this study, we introduced TransHLA, a pre-trained language model-based deep learning framework for predicting epitopes presented by both HLA-I and HLA-II. TransHLA uses a large language model to extract structural and text features of candidate sequences and then processes these features using Convolutional Neural Networks (CNN) and Transformer modules. Experimental results on the benchmark dataset show that TransHLA outperforms cutting edge sequence classification models in predicting both HLA-I and HLA-II binding epitopes.

In comparison to traditional epitope-HLA binding methods, which rely on both epitope and HLA allele information, TransHLA allows users to perform epitope screening without requiring HLA alleles as input. While HLA typing can be accurately performed using RNA-seq or WES data in personalized medicine contexts, TransHLA is designed to streamline allele-specific binding predictions during the screening of immunogenic epitopes, serving as a preliminary step for the widely used tools that focus on HLA-epitope binding affinity. When discovering new epitopes, we observed that binding affinity models tend to more easily misclassify ordinary peptide sequences as epitopes as the number of input alleles increases. This phenomenon arises because, despite binding affinity models being trained on benchmark datasets, the inclusion of multiple alleles introduces increased sequence diversity and variability in binding motifs or features, leading to higher uncertainty and reduced prediction accuracy. Additionally, for certain alleles with a disproportionately low number of epitopes available for training or other limitations, the insufficient training further exacerbates this issue. For example, in netMHCpan4.1, the allele HLA-A03:02 has only 13 epitopes available for training, resulting in an AUC of just 0.6331. As a result of these challenges, these models often misidentify non-presented negative samples as potential epitopes, which increases the experimental burden and adds additional costs for epitope discovery. This limitation is particularly problematic in scenarios such as selecting antigens for a population-wide vaccination, where the population contains many different alleles. In such cases, the variability across alleles further complicates the ability of binding affinity models to reliably identify epitopes. From Table 3 and Table 4, it is evident that binding affinity prediction software achieves significantly lower specificity in epitope detection tasks compared to TransHLA, further highlighting the limitations of these models. TransHLA effectively addresses this issue by adopting a novel training strategy: combining all epitopes into a single class as positive samples and using a 1:1 ratio of negative samples for training. This approach allows the predictor to learn the unified features of epitopes, enabling it to better distinguish epitopes from ordinary peptide sequences. By leveraging this strategy, TransHLA not only retains most potential epitopes while filtering out non-presented negatives, significantly reducing the need for extensive chemical experiments and lowering costs, but also proves particularly advantageous in tasks like population-wide antigen selection by improving the robustness and reliability of epitope detection.

In conclusion, TransHLA serves as a powerful complementary tool that expedite the precise screening of epitopes. TransHLA efficiently filters out non-epitope sequences and achieves higher accuracy compared to conventional methods. In a general Neoeptope dataset verified by TCR experiments, TransHLA achieves an accuracy of 90.65% for HLA-I epitopes.

## Availability of source code and requirements

- Project name: TransHLA
- Project home page: https://github.com/SkywalkerLuke/TransHLA
- RRID: SCR _ 026171
- BioTools ID: biotools: transhla
- Operating system(s): Platform independent
- Programming language: Python
- Other requirements: Python 3.9 or higher, pytorch 2.0 or higher
- License: MIT license

## Data Availability

The datasets used in this paper are collected from IEDB, CEDAR, VDJdb, ImmunoCode, dbpepneo2.0 and NEPdb [20, 21, 22, 23, 24, 25]. An archival copy of the code and supporting is available via the GigaScience database, GigaDB [55]. We also submit the training data and test data along with our source code of TransHLA. The alleles used by the software for predicting HLA-binding affinity are also provided in GigaDB. DOME-ML (Data, Optimization, Model and Evaluation in Machine Learning) annotations are available in the DOME registry via accession peuywb6nkx[56].

## Author’s Contributions

TCL and XYW designed the study and algorithm, implemented the code, and wrote the manuscript. SCL supervised the project, designed the initial framework, and revised the manuscript. NW implemented the tests and discussed the project. MZH discussed the project. All authors read and approved the final manuscript.

## References

1. Chaffey N, Alberts, B., Johnson, A., Lewis, J., Raff, M., Roberts, K. and Walter, P. Molecular biology of the cell. 4th edn. Oxford University Press; 2003.

2. Murphy K, Weaver C. Janeway’s immunobiology. Garland science; 2016.

3. Abbas A, Lichtman A, Pillai S. Cellular and molecular immunology E-book. Elsevier Health Sciences; 2014.

4. Shao XM, Bhattacharya R, Huang J, Sivakumar IA, Tokheim C, Zheng L, et al. High-throughput prediction of MHC class I and II neoantigens with MHCnuggets. Cancer immunology research 2020;8(3):396-408.

5. O’Donnell TJ, Rubinsteyn A, Laserson U. MHCflurry 2.0: improved pan-allele prediction of MHC class I-presented peptides by incorporating antigen processing. Cell systems 2020;11(1):42-48.

6. Liu Z, Jin J, Cui Y, Xiong Z, Nasiri A, Zhao Y, et al. DeepSeq-PanII: an interpretable recurrent neural network model with attention mechanism for peptide-HLA class II binding prediction. IEEE/ACM Transactions on Computational Biology and Bioinformatics 2021;19(4):2188-2196.

7. Nguyen AT, Szeto C, Gras S. The pockets guide to HLA class I molecules. Biochemical Society Transactions 2021;49(5):2319- 2331.

8. Mei S, Li F, Xiang D, Ayala R, Faridi P, Webb GI, et al. Anthem: a user customised tool for fast and accurate prediction of binding between peptides and HLA class I molecules. Briefings in Bioinformatics 2021;22(5):bbaa415.

9. Reynisson B, Alvarez B, Paul S, Peters B, Nielsen M. NetMHCpan-4.1 and NetMHCIIpan-4.0: improved predictions of MHC antigen presentation by concurrent motif deconvolution and integration of MS MHC eluted ligand data. Nucleic acids research 2020;48(W1):W449-W454.

10. Chu Y, Zhang Y, Wang Q, Zhang L, Wang X, Wang Y, et al. A transformer-based model to predict peptide-HLA class I binding and optimize mutated peptides for vaccine design. Nature Machine Intelligence 2022;4(3):300-311.

11. Tadros DM, Racle J, Gfeller D. Predicting MHC-I ligands across alleles and species: How far can we go? bioRxiv 2024;p. 2024- 05.

12. Racle J, Guillaume P, Schmidt J, Michaux J, Larabi A, Lau K, et al. Machine learning predictions of MHC-II specificities reveal alternative binding mode of class II epitopes. Immunity 2023;56(6):1359-1375.

13. Vaswani A, Shazeer N, Parmar N, Uszkoreit J, Jones L, Gomez AN, et al. Attention is all you need. Advances in neural information processing systems 2017;30.

14. He K, Zhang X, Ren S, Sun J. Deep residual learning for image recognition. In: Proceedings of the IEEE conference on computer vision and pattern recognition; 2016. p. 770-778.

15. Lin Z, Akin H, Rao R, Hie B, Zhu Z, Lu W, et al. Evolutionary-scale prediction of atomic-level protein structure with a language model. Science 2023;379(6637):1123-1130.

16. Kim Y. Convolutional neural networks for sentence classification. arXiv preprint arXiv:14085882 2014;.

17. Lai S, Xu L, Liu K, Zhao J. Recurrent convolutional neural networks for text classification. In: Proceedings of the AAAI conference on artificial intelligence, vol. 29; 2015..

18. Johnson R, Zhang T. Deep pyramid convolutional neural networks for text categorization. In: Proceedings of the 55th Annual Meeting of the Association for Computational Linguistics (Volume 1: Long Papers); 2017. p. 562-570.

19. Zhou P, Shi W, Tian J, Qi Z, Li B, Hao H, et al. Attention-based bidirectional long short-term memory networks for relation classification. In: Proceedings of the 54th annual meeting of the association for computational linguistics (volume 2: Short papers); 2016. p. 207-212.

20. Vita R, Mahajan S, Overton JA, Dhanda SK, Martini S, Cantrell JR, et al. The immune epitope database (IEDB): 2018 update. Nucleic acids research 2019;47(D1):D339-D343.

21. Koşaloğlu-Yalçın Z, Blazeska N, Vita R, Carter H, Nielsen M, Schoenberger S, et al. The cancer epitope database and analysis resource (CEDAR). Nucleic Acids Research 2023;51(D1):D845- D852.

22. Shugay M, Bagaev DV, Zvyagin IV, Vroomans RM, Crawford JC, Dolton G, et al. VDJdb: a curated database of T-cell receptor sequences with known antigen specificity. Nucleic acids research 2018;46(D1):D419-D427.

23. Nolan S, Vignali M, Klinger M, Dines JN, Kaplan IM, Svejnoha E, et al. A large-scale database of T-cell receptor beta (TCRβ) sequences and binding associations from natural and synthetic exposure to SARS-CoV-2. Research square 2020;.

24. Lu M, Xu L, Jian X, Tan X, Zhao J, Liu Z, et al. dbPepNeo2. 0: A database for human tumor neoantigen peptides from mass spectrometry and TCR recognition. Frontiers in Immunology 2022;13:855976.

25. Xia J, Bai P, Fan W, Li Q, Li Y, Wang D, et al. NEPdb: a database of T-cell experimentally-validated neoantigens and pan-cancer predicted neoepitopes for cancer immunotherapy. Frontiers in Immunology 2021;12:644637.

26. Buchfink B, Xie C, Huson DH. Fast and sensitive protein alignment using DIAMOND. Nature methods 2015;12(1):59-60.

27. Pruitt KD, Tatusova T, Maglott DR. NCBI reference sequences (RefSeq): a curated non-redundant sequence database of genomes, transcripts and proteins. Nucleic acids research 2007;35(Suppl_1):D61-D65.

28. Fu L, Niu B, Zhu Z, Wu S, Li W. CD-HIT: accelerated for clustering the next-generation sequencing data. Bioinformatics 2012;28(23):3150-3152.

29. Chen B, Cheng X, Li P, Geng Ya, Gong J, Li S, et al. xTrimoPGLM: unified 100B-scale pre-trained transformer for deciphering the language of protein. arXiv preprint 240106199 2024;.

30. Elnaggar A, Heinzinger M, Dallago C, Rehawi G, Yu W, Jones L, et al. ProtTrans: Towards Cracking the Language of Lifes Code Through Self-Supervised Deep Learning and High Performance Computing. IEEE Transactions on Pattern Analysis and Machine Intelligence 2021;p. 1-1.

31. Brown TB, Mann B, Ryder N, Subbiah M, Kaplan J, Dhariwal P, et al. Language Models are Few-Shot Learners. Advances in Neural Information Processing Systems 2020;33:1877-1901.

32. Du Z, Ding X, Xu Y, Li Y. UniDL4BioPep: a universal deep learning architecture for binary classification in peptide bioactivity. Briefings in Bioinformatics 2023;24(3):bbad135.

33. Xu Z, Zhong H, He B, Wang X, Lu T. PTransIPs: Identification of Phosphorylation Sites Enhanced by Protein PLM Embeddings. IEEE Journal of Biomedical and Health Informatics 2024;.

34. Jumper J, Evans R, Pritzel A, Green T, Figurnov M, Ronneberger O, et al. Highly accurate protein structure prediction with AlphaFold. Nature 2021;596(7873):583-589.

35. Mirdita M, Schütze K, Moriwaki Y, Heo L, Ovchinnikov S, Steinegger M. ColabFold: making protein folding accessible to all. Nature methods 2022;19(6):679-682.

36. Devlin J, Chang MW, Lee K, Toutanova K. Bert: Pre-training of deep bidirectional transformers for language understanding. arXiv preprint 181004805 2018;.

37. Boudiaf M, Masud ZI, Rony J, Dolz J, Piantanida P, Ayed IB, Transductive Information Maximization For Few-Shot Learning; 2020.

38. Racle J, Michaux J, Rockinger GA, Arnaud M, Bobisse S, Chong C, et al. Robust prediction of HLA class II epitopes by deep motif deconvolution of immunopeptidomes. Nature biotechnology 2019;37(11):1283-1286.

39. Nagler A, Kalaora S, Barbolin C, Gangaev A, Ketelaars SL, Alon M, et al. Identification of presented SARS-CoV-2 HLA class I and HLA class II peptides using HLA peptidomics. Cell Reports 2021;35(13).

40. Yang Y, Wei Z, Cia G, Song X, Pucci F, Rooman M, et al. MHCII-peptide presentation: an assessment of the state-of-the-art prediction methods. Frontiers in Immunology 2024;15:1293706.

41. Maćkiewicz A, Ratajczak W. Principal components analysis (PCA). Computers & Geosciences 1993;19(3):303-342.

42. Westhof E, Altschuh D, Moras D, Bloomer A, Mondragon A, Klug A, et al. Correlation between segmental mobility and the location of antigenic determinants in proteins. Nature 1984;311(5982):123-126.

43. Kim DG, Choi Y, Kim HS. Epitopes of protein binders are related to the structural flexibility of a target protein surface. Journal of Chemical Information and Modeling 2021;61(4):2099-2107.

44. Klatt MG, Mack KN, Bai Y, Aretz ZE, Nathan LI, Mun SS, et al. Solving an MHC allele-specific bias in the reported immunopeptidome. JCI insight 2020;5(19).

45. Cock PJ, Antao T, Chang JT, Chapman BA, Cox CJ, Dalke A, et al. Biopython: freely available Python tools for computational molecular biology and bioinformatics. Bioinformatics 2009;25(11):1422.

46. Vihinen M, Torkkila E, Riikonen P. Accuracy of protein flexibility predictions. Proteins: Structure, Function, and Bioinformatics 1994;19(2):141-149.

47. Chen T, Guestrin C. Xgboost: A scalable tree boosting system. In: Proceedings of the 22nd acm sigkdd international conference on knowledge discovery and data mining; 2016. p. 785-794.

48. Mariño Pérez L, Ielasi FS, Bessa LM, Maurin D, Kragelj J, Blackledge M, et al. Visualizing protein breathing motions associated with aromatic ring flipping. Nature 2022;602(7898):695-700.

49. Anjana R, Vaishnavi MK, Sherlin D, Kumar SP, Naveen K, Kanth PS, et al. Aromatic-aromatic interactions in structures of proteins and protein-DNA complexes: a study based on orientation and distance. Bioinformation 2012;8(24):1220.

50. Baker E. Hydrogen bonding in biological macromolecules 2012;.

51. Grimaldi J, Radhakrishna M, Kumar SK, Belfort G. Stability of proteins on hydrophilic surfaces. Langmuir 2015;31(3):1005- 1010.

52. Drelich J, Chibowski E, Meng DD, Terpilowski K. Hydrophilic and superhydrophilic surfaces and materials. Soft Matter 2011;7(21):9804-9828.

53. Perticaroli S, Nickels JD, Ehlers G, O’Neill H, Zhang Q, Sokolov AP. Secondary structure and rigidity in model proteins. Soft Matter 2013;9(40):9548-9556.

54. Mamonova TB, Glyakina AV, Galzitskaya OV, Kurnikova MG. Stability and rigidity/flexibility—Two sides of the same coin? Biochimica et Biophysica Acta (BBA)-Proteins and Proteomics 2013;1834(5):854-866.

55. Tianchi L, Xueying W, Wan N, Huo M, Shuaicheng L. Supporting data for “TransHLA: A Hybrid Transformer Model for HLA-Presented Epitope Detection”. GigaDB 2024; https://urldefense.com/v3/__https://doi.org/10.5524/102633__;!!KjDnqvtInNPT!i0agEmhs5f2c-6xn78vf1UHeQDc7MIvRJjXtTKhowEM818-6OF2FSrgCa2kiVF4O5NMkHRJhDcvGVR-Dmyvs$.

56. Tianchi L, Xueying W, Wan N, Huo M, Shuaicheng L, TransHLA: A Hybrid Transformer Model for HLA-Presented Epitope Detection; 2024. https://urldefense.com/v3/__https://registry.dome-ml.org/review/peuywb6nkx__;!!KjDnqvtInNPT!i0agEmhs5f2c-6xn78vf1UHeQDc7MIvRJjXtTKhowEM818-6OF2FSrgCa2kiVF4O5NMkHRJhDcvGVW88ZMPB$. [DOME-ML Annotations].

