## supplementary file for "TransHLA: A Hybrid Transformer Model for HLA-Presented Epitope Detection"

### Supplementary file 1

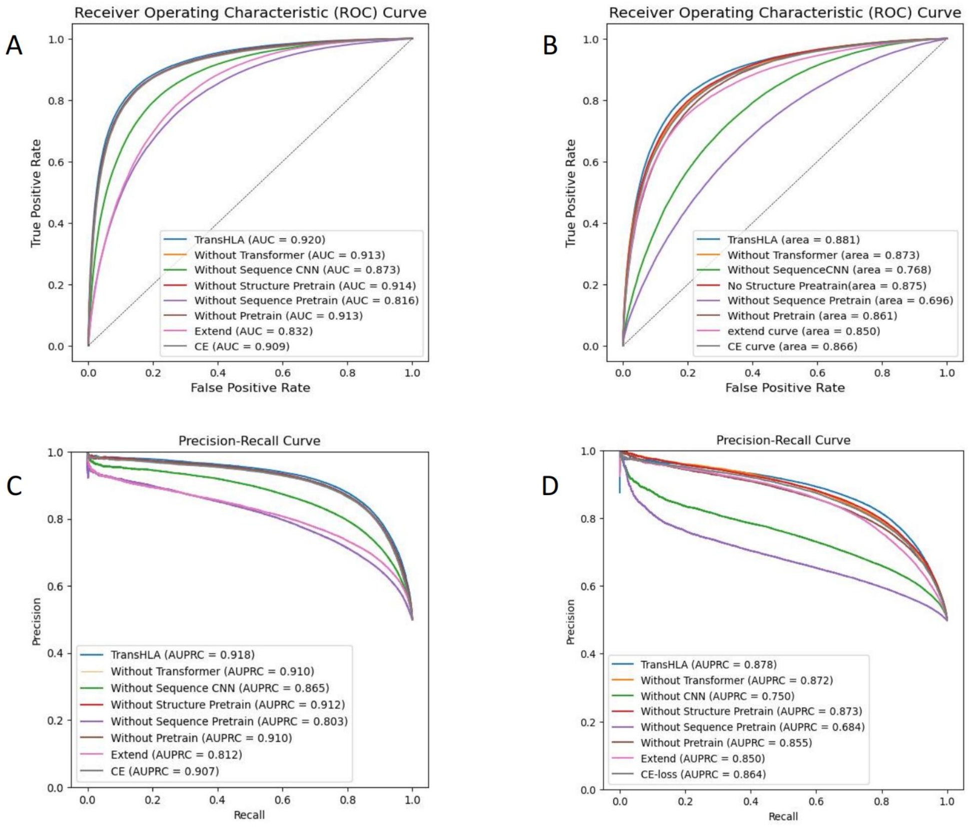


**Supplementary Figure 1.** The figure presents an ablation study for our predictive model in different modules, showing how performance metrics—AUROC and AUPRC—are affected by the removal of specific modules for both HLA-I (Subfigures A and C) and HLA-II (Subfigures B and D) classes. Changes in these metrics underscore the contribution of each component to the model's accuracy in epitope prediction, providing insight into the model's architecture and the pivotal elements for its effectiveness across HLA classes.

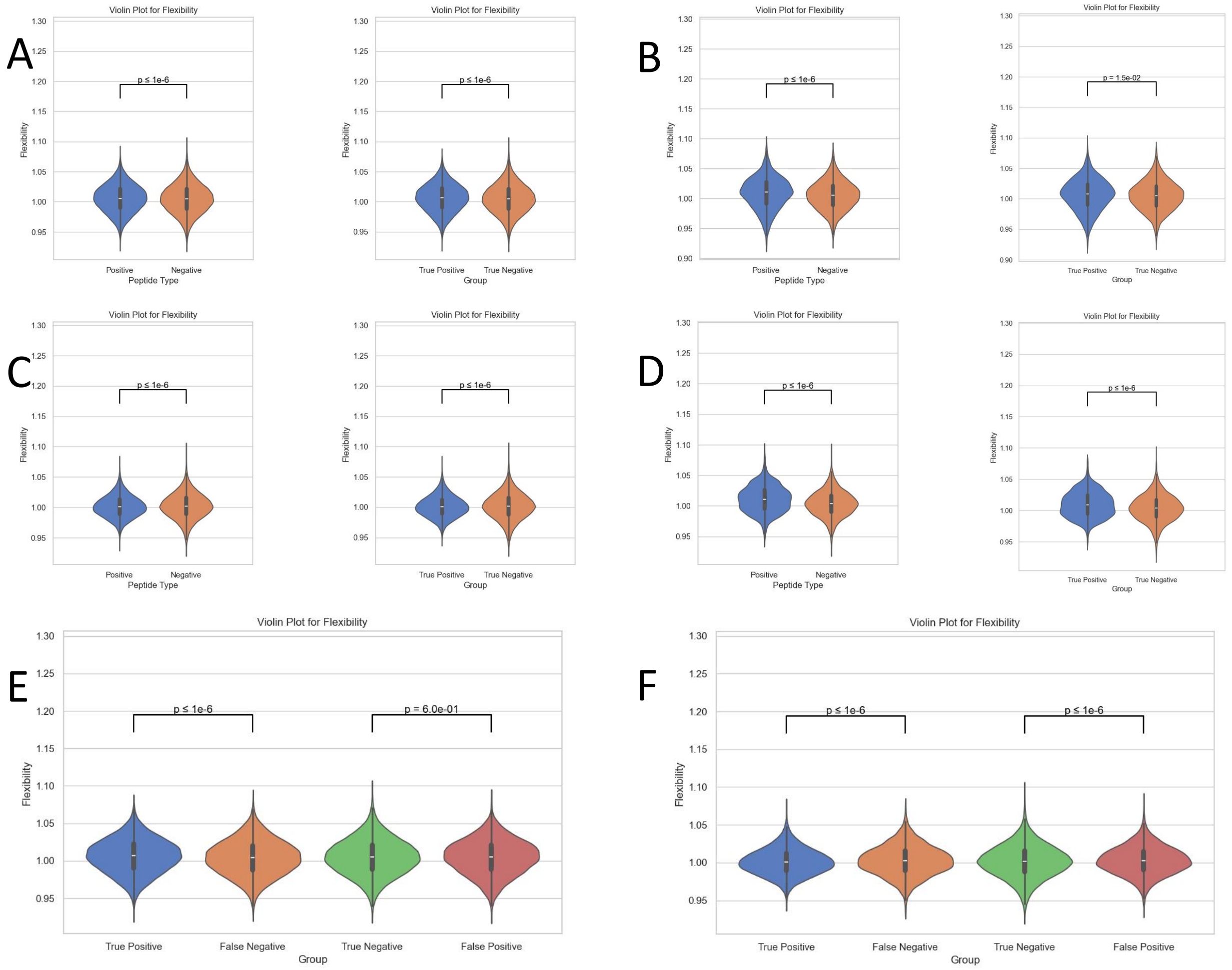


**Supplementary Figure 2.** This comprehensive figure presents a series of violin plots illustrating the 'Flexibility' chemical property of peptides across various sample subsets for HLA-I and HLA-II molecules. Subfigure (A) delineates the Flexibility distribution in independent test samples for HLA-I, separated into positive and negative samples, with each subgroup's statistical significance assessed via t-tests and annotated with corresponding p-values. Subfigure (C) mirrors this setup for HLA-II independent test samples, highlighting the comparative Flexibility distributions. The external dataset distributions for HLA-I and HLA-II are respectively showcased in subfigures (B) and (D), emphasizing the metric's external validity. Subfigures (E) and (F) delve deeper, contrasting the Flexibility of true positives and false negatives against true negatives and false positives within HLA-I and HLA-II datasets, respectively.

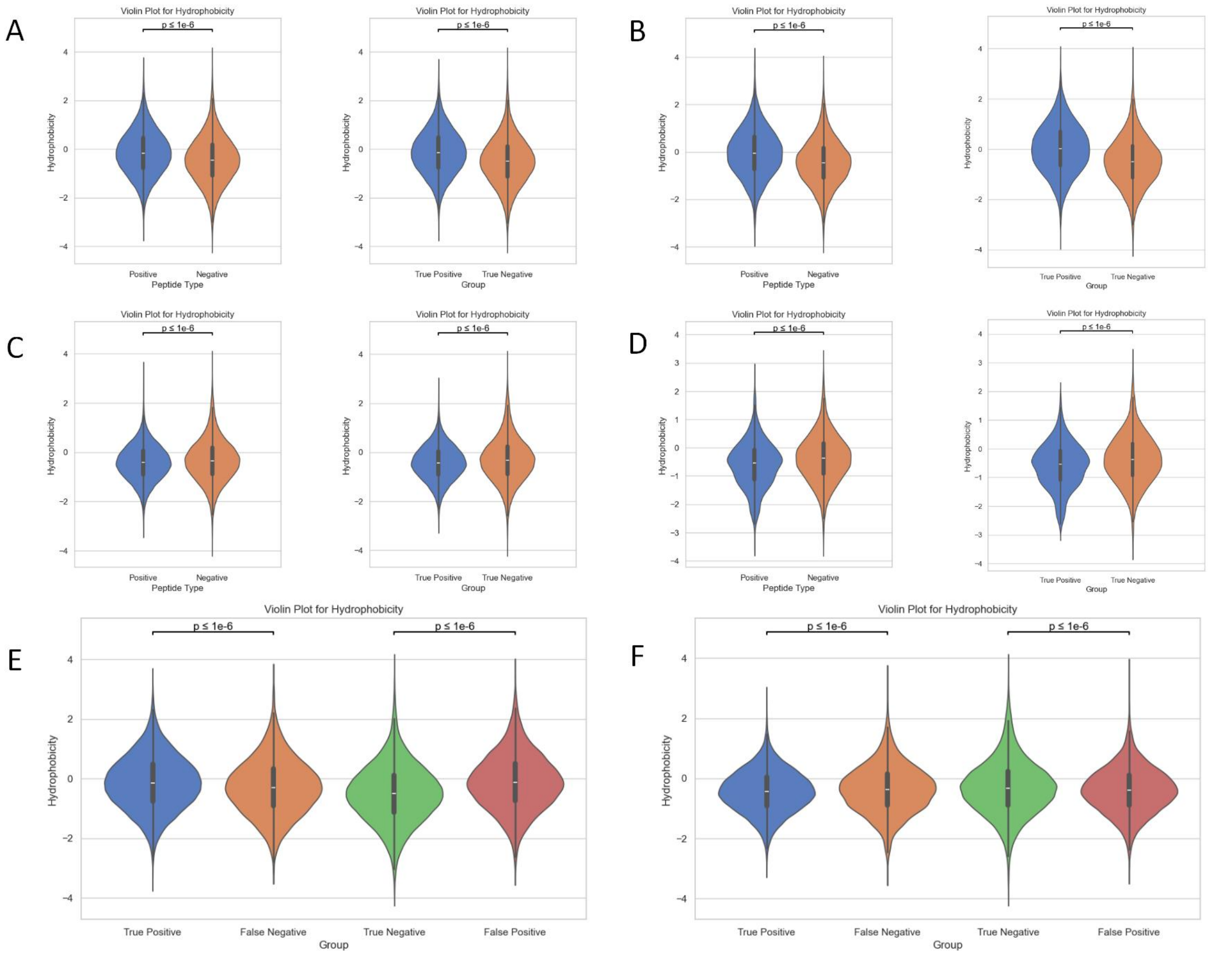


**Supplementary Figure 3.** This comprehensive figure presents a series of violin plots illustrating the 'Hydrophobicity' chemical property of peptides across various sample subsets for HLA-I and HLA-II molecules. Subfigure (A) delineates the Hydrophobicity distribution in independent test samples for HLA-I, separated into positive and negative samples, with each subgroup's statistical significance assessed via t-tests and annotated with corresponding p-values. Subfigure (C) mirrors this setup for HLA-II independent test samples, highlighting the comparative Hydrophobicity distributions. The external dataset distributions for HLA-I and HLA-II are respectively showcased in subfigures (B) and (D), emphasizing the metric's external validity. Subfigures (E) and (F) delve deeper, contrasting the Hydrophobicity of true positives and false negatives against true negatives and false positives within HLA-I and HLA-II datasets, respectively.

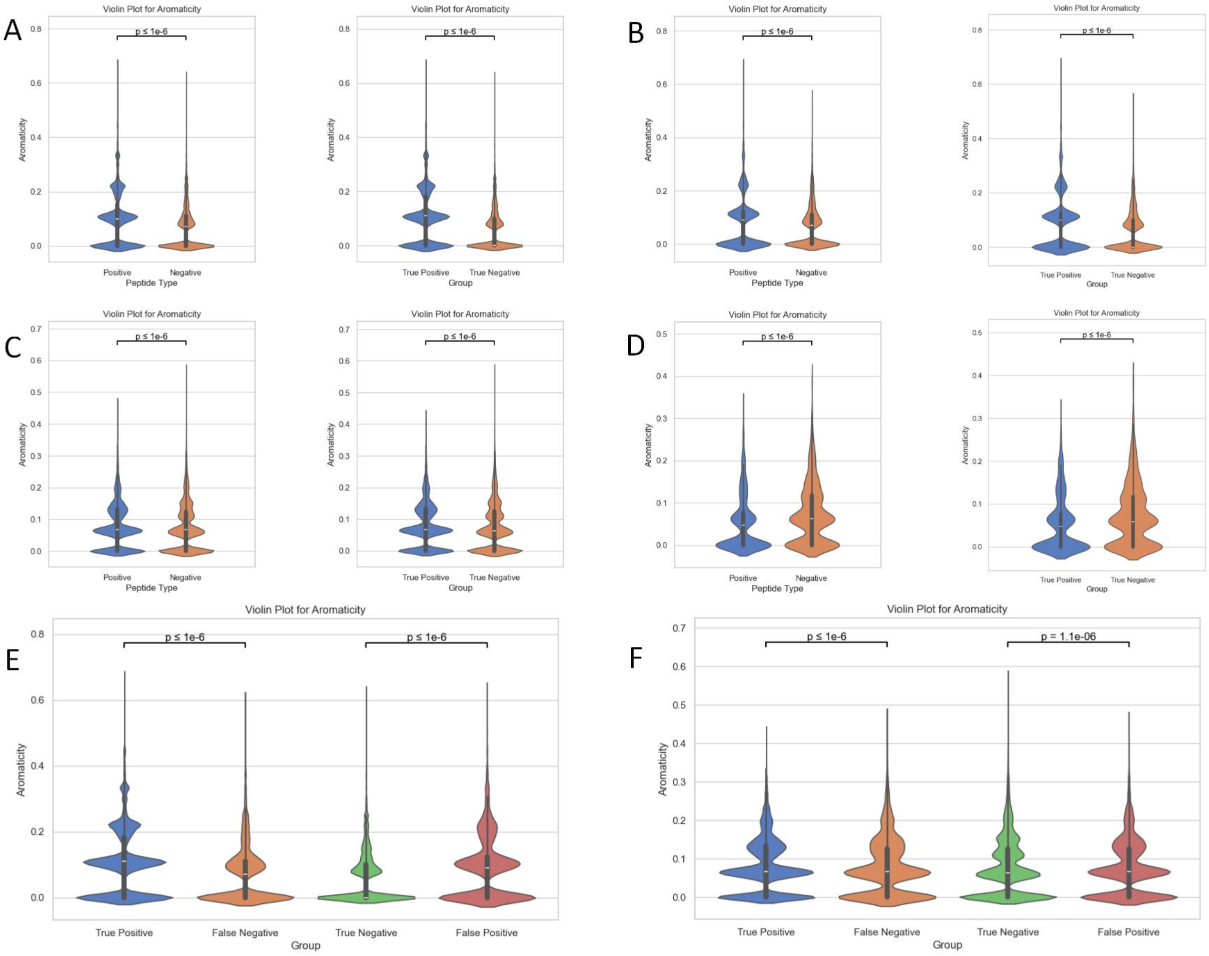


**Supplementary Figure 4.** This comprehensive figure presents a series of violin plots illustrating the 'Aromaticity' chemical property of peptides across various sample subsets for HLA-I and HLA-II molecules. Subfigure (A) delineates the Aromaticity distribution in independent test samples for HLA-I, separated into positive and negative samples, with each subgroup's statistical significance assessed via t-tests and annotated with corresponding p-values. Subfigure (C) mirrors this setup for HLA-II independent test samples, highlighting the comparative Aromaticity distributions. The external dataset distributions for HLA-I and HLA-II are respectively showcased in subfigures (B) and (D), emphasizing the metric's external validity. Subfigures (E) and (F) delve deeper, contrasting the Aromaticity of true positives and false negatives against true negatives and false positives within HLA-I and HLA-II datasets, respectively.

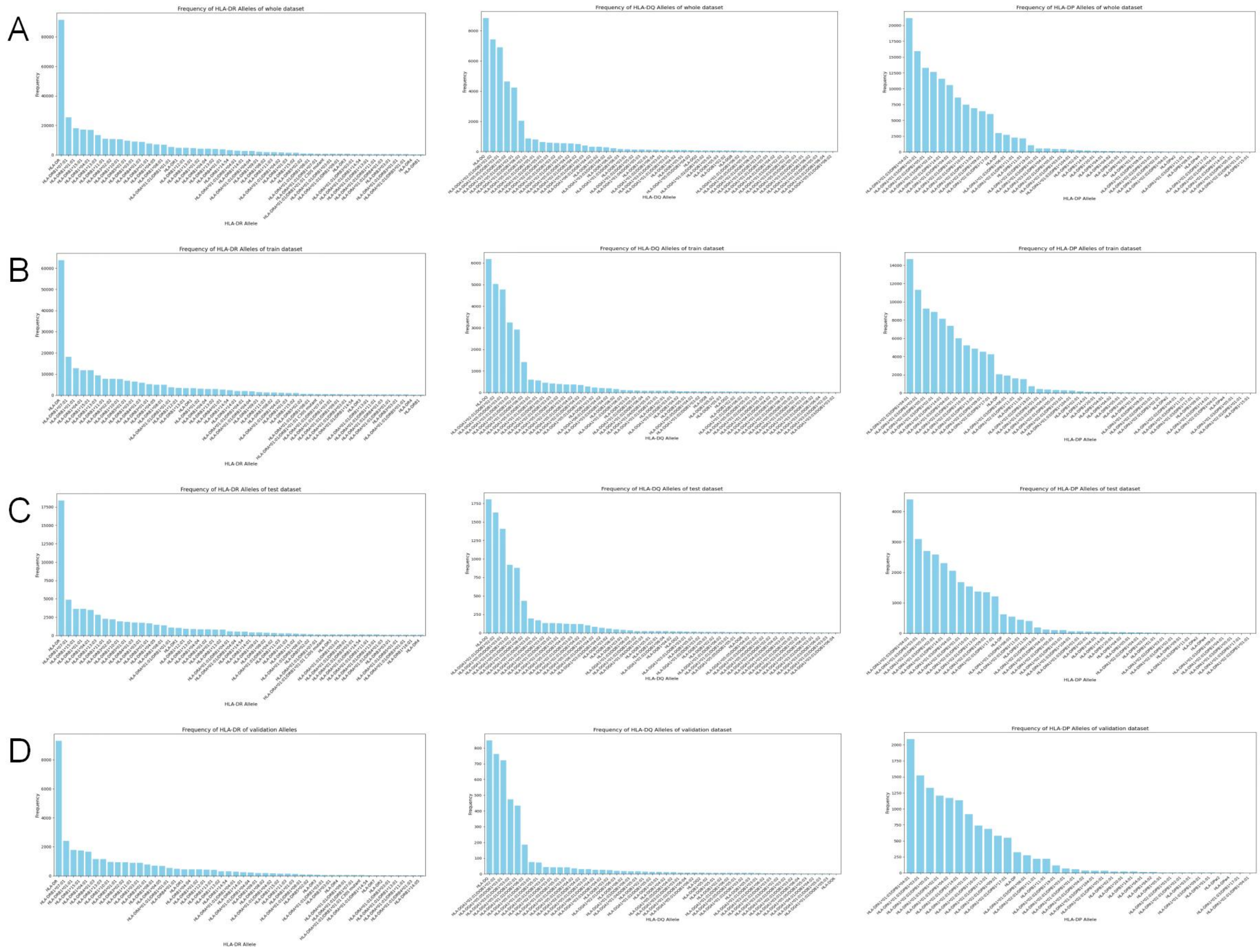


**Supplementary Figure 5.** This figure illustrates the experimental distribution of major HLA-II alleles in the IEDB. We plotted the alleles by type, where subfigure A shows the distribution for all data, B for the training dataset, C for the test dataset, and D for the validation dataset. Left represents HLA-DR alleles, middle represents HLA-DQ alleles, and right represents HLA-DP alleles.

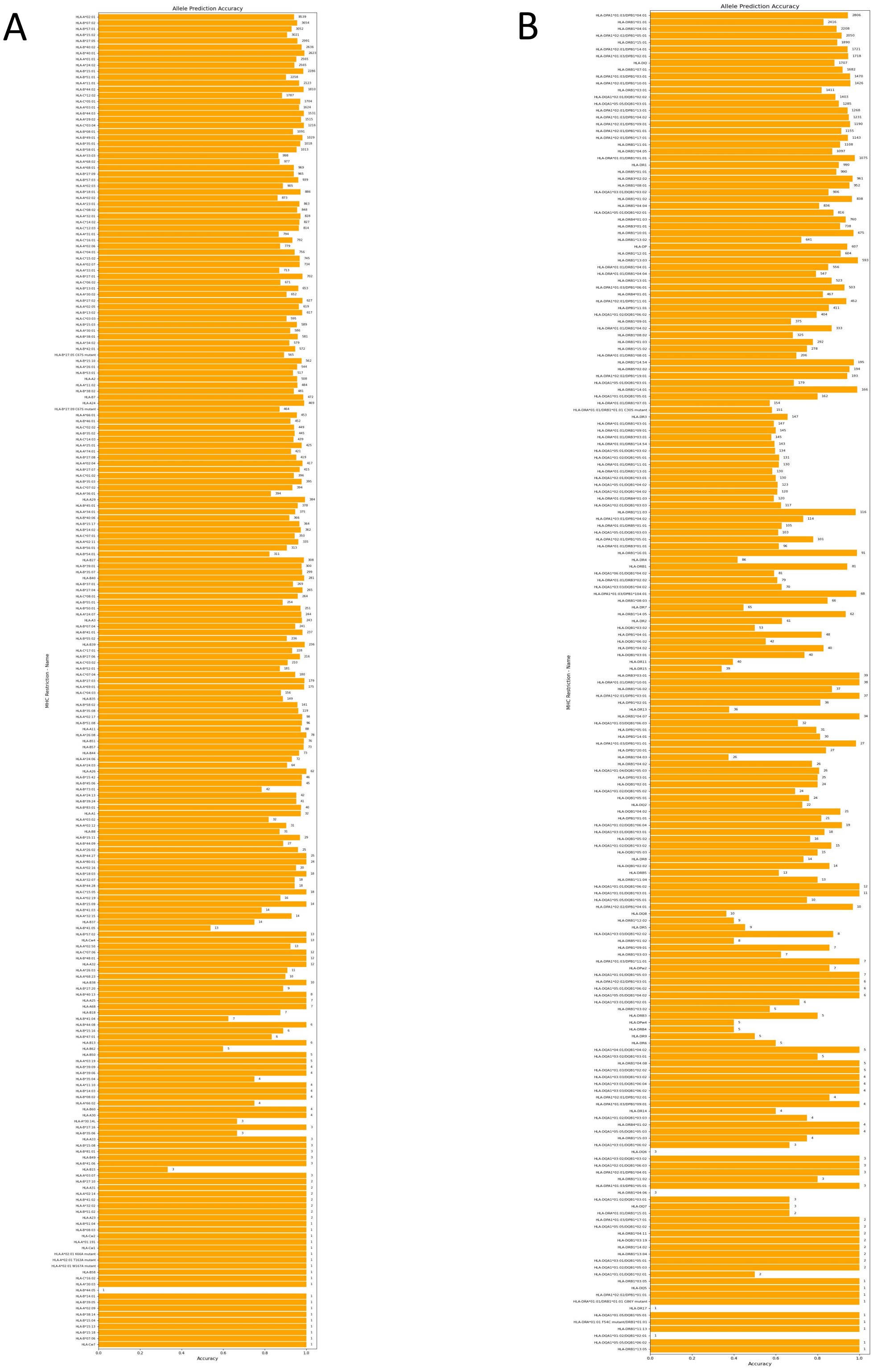

**Supplementary Figure 6.** This figure illustrates the prediction results of TransHLA on the test set for different HLA alleles. Panel A shows the results corresponding to HLA-I alleles, while Panel B displays those for HLA-II alleles. The number of epitopes corresponding to each allele is annotated to the right of each bar.

| Feature | HLA I Importance | HLA II Importance |
| --- | --- | --- |
| Helix Content | 0.315736 | 0.166237 |
| Aromaticity | 0.257780 | 0.218409 |
| Flexibility | 0.211219 | 0.046253 |
| Sheet Content | 0.063786 | 0.091180 |
| Isoelectric Point | 0.043059 | 0.111911 |
| Mean-Charge-at-PH-7 | 0.034550 | 0.089355 |
| Coil Content | 0.033767 | 0.048550 |
| Hydrophobicity | 0.016858 | 0.129952 |
| Mean Molecular Weight | 0.013744 | 0.052652 |
| Instability Index | 0.009502 | 0.045500 |

**Supplementary Table 1: The importance of different Features in XGboost model**

| Type | ACC(%) | F1(%) | Recall(%) | MCC | Precision(%) | Specificity (%) |
| --- | --- | --- | --- | --- | --- | --- |
| HLA-I | 84.75 | 67.73 | 80.06 | 0.59 | 58.70 | 86.11 |
| HLA-I | 71.14 | 45.08 | 59.24 | 0.28 | 36.39 | 74.12 |

**Supplementary Table 2: TransHLA's performance in Four-time Negative samples dataset**

| HLA I Type | Epitopes | Prediction Probability |
| --- | --- | --- |
| HLA I | ALWGFFPVL | 0.9669 |
|  | LDTNADKQLSF | 0.8441 |
|  | WQQGLRVSF | 0.9774 |
|  | ILDTAGKEEY | 0.9930 |
| HLA II | PKYVKQNTLKLAT | 0.9993 |
|  | ISTNIRQAGVQYSRA | 0.9892 |

**Supplementary Table 3: TransHLA's performance of allele unseen epitopes in test dataset**

| HLA Restriction | Epitope |
|---|---|
| HLA-A*02:01 T163A mutant | ALWGFFPVL |
| HLA-A*02:01 W167A mutant | ALWGFFPVL |
| HLA-A*02:01 K66A mutant | ALWGFFPVL |
| HLA-Cw2 | LDTNADKQLSF |
| HLA-B*15:04 | WQQGLRVSF |
| HLA-A*01:191 | ILDTAGKEEY |
| HLA-DRB1*13:04 | PKYVKQNTLKLAT |
| HLA-DRB1*13:05 | PKYVKQNTLKLAT |
| HLA-DRA*01:01/DRB1*01:01 G86Y mutant | PKYVKQNTLKLAT |
| HLA-DRB1*13:04 | ISTNIRQAGVQYSRA |

**Supplementary Table 4: The unseen alleles epitopes in test data and their corresponding alleles**

#### TransHLA Architecture Hyperparameters

The TransHLA model is specifically designed for applications in immunogenetics, focusing on the analysis of HLA (Human Leukocyte Antigen) sequences. It incorporates a hybrid architecture featuring both Transformer and Convolutional Neural Network (CNN) components. Below are detailed descriptions of the model components and their respective hyperparameters:

**Model Components**

- **Pretrained ESM-2 Model:** Uses the esm2_t33_650M_UR50D model for the initial embedding of sequences.
- **CNN Layers for Region and Structure Embedding:**
  - **Layer 1** (region_cnn1) and **Layer 2** (region_cnn2): Each serves different embedding purposes tailored to specific features of the input sequences.
- **Transformer Encoder:**
  - **Number of Layers** (n_layers): Determined by model-specific parameters, accommodating varying depths of processing.
  - **Number of Attention Heads** (n_head): Configured to optimize attention mechanisms across different parts of the input sequence.
  - **Model Dimension** (d_model): Defines the size of feature vectors within the transformer, influencing the capacity to handle information complexity.
  - **Feedforward Dimension** (d_ff): Specifies the size of the internal feed-forward networks, crucial for processing layers of features.
  - **Batch Normalization:** Implemented after certain layers to enhance model stability during training.
  - **Max Pooling and Padding:** Employed strategically within CNN layers to effectively reduce dimensionality and adjust tensor sizes.

**Hyperparameters**

- **Maximum Input Length** (max_len): 512 - Sets the limit on the length of sequences the model can process at once.
- **CNN Padding Index** (cnn_padding_index): 0 - Indicates the padding value used in CNN layers to maintain consistency in tensor dimensions.
- **CNN Number of Channels** (cnn_num_channel): 256 - Specifies the output channels of the CNN layers, directly influencing the breadth of feature detection.
- **Region Embedding Size** (region_embedding_size): Typically 3 filters that capture local contextual information within the sequence.
- **CNN Kernel Size** (cnn_kernel_size): 3 - Determines the extent of each convolution operation, affecting how the input data is processed.
- **CNN Padding Size** (cnn_padding_size): 1 - Ensures that the convolution output maintains appropriate dimensions.
- **CNN Stride** (cnn_stride): 1 - Controls the movement of the convolutional filters across the input field.
- **Pooling Size** (pooling_size): 2 - Defines the window size for max pooling, reducing spatial dimensions while preserving the most significant features.

**Forward Pass Description**

- **Input Processing:** Starts by embedding sequences using a pretrained ESM model, from which both sequence and structural representations are derived.

- **Transformer Encoding:** Advances the embeddings through a series of Transformer encoder layers, processing the complex dependencies within the data.
- **CNN Processing:** Applies a sequence of CNN layers to both sequence and structure-derived embeddings, enhancing feature extraction through residual connections and pooling techniques.
- **Feature Integration and Classification:**
    - Integrates outputs from both the Transformer and CNN pathways.
    - Propels the integrated features through dense layers for further refinement.
    - Concludes with a softmax layer to output classification probabilities, providing predictions on HLA types or related immunological responses.

This architecture is meticulously crafted to integrate intricate sequence and structural information, ensuring precise predictions in immunogenetics applications.

#### Deep Pyramid Convolutional Neural Network (DPCNN) Architecture Hyperparameters

The DPCNN model is designed for efficient text classification leveraging deep pyramid structures. Below we provide the hyperparameters used in the DPCNN architecture as described in the manuscript.

**Embedding Layer:**

- **Vocabulary Size (num_vocab):** 1000 words
- **Embedding Dimension (embedding_dim):** 256
- **Padding Index (padding_index):** 0 (zero-padding for alignment)
- Region Embedding:
- **Region Embedding Size:** Size of each region for initial feature extraction

**Convolutional Layers:**

- **Number of Channels (cnn_num_channel):** 256 channels for convolutional operations
- **Kernel Size (cnn_kernel_size):** 3 (width of the convolutional kernel)
- **Padding Size (cnn_padding_size):** 1 (padding applied to each side of the input)
- **Stride (cnn_stride):** 1 (stride of the convolutional operation)

Pooling Layer:

- **Pooling Size (pooling_size):** 2 (factor by which to downsample the input)

Output Layer:

- **Number of Classes (num_classes):** 2 (binary classification)

The model incorporates region embedding followed by two sets of convolutional operations with ReLU activation and constant padding. The output from the convolutional layers passes through max pooling before being flattened and passed to a fully connected layer for classification. The architecture employs residual connections and is optimized to handle varying lengths of text input with a focus on extracting hierarchical features for text representation.

#### Recurrent Neural Network with Attention (RNN_ATTs) Architecture Hyperparameters

The RNN_ATTs model incorporates an attention mechanism over a recurrent neural network for sequence modeling tasks. The following hyperparameters define the architecture:

- **Vocabulary Size:** 40
- **Embedding Dimension:** 256
- **Hidden Dimension (LSTM):** 128
- **Number of LSTM Layers:** 2
  **Bidirectionality:** Enabled (True)
  **Dropout Rate:** 0.2
- **Padding Index:** 0
- **Second Hidden Layer Size:** 64
- **Output Dimension:** 2 (binary classification)

The model uses an LSTM layer with tanh activation for sequence processing, followed by an attention mechanism that assigns weights to the LSTM outputs. The attention-weighted outputs are then passed through a fully connected layer with ReLU activation before reaching the final classification layer.

#### TextCNN Architecture Hyperparameters

The TextCNN model applies convolutional neural networks to text classification tasks. Here are the key hyperparameters:

- **Vocabulary Size:** 40 tokens
- **Embedding Dimension:** 128
- **Window Sizes:** [2, 4, 3] (corresponding to the size of the convolutional filters)
- **Maximum Sequence Length:** 21
- **Feature Size (Number of Convolutional Filters):** 256 per window size
- **Number of Classes (n_class):** 2 (binary classification)
- **Dropout Rate:** 0.4

The architecture consists of an embedding layer, followed by parallel convolutional layers with varying window sizes for feature extraction, and a max-pooling operation. The pooled features are concatenated and passed through a dropout layer before a fully connected layer for classification.

#### TextRCNN (Text Recurrent Convolutional Neural Network) Architecture Hyperparameters

The TextRCNN model combines recurrent neural network (RNN) with convolutional neural network (CNN) concepts for text classification. Here are the essential hyperparameters and components of the architecture:

- **Vocabulary Size:** 40 tokens
- **Embedding Dimension:** 128
- **Hidden Size (LSTM):** 50

- **Number of Labels (Output Dimension):** 2 (binary classification)
- **Dropout Rate:** 0.5

The architecture employs the following layers and operations:

- An embedding layer that maps tokens to vectors of the specified embedding dimension.
- A bidirectional LSTM layer for processing sequences, capturing both forward and backward context.
- A custom GlobalMaxPool1d layer that applies global max pooling over the time dimension of the LSTM output.
- Two linear layers with ReLU activation after the first linear transformation. The first linear layer expands the concatenated LSTM outputs and embedding vectors to a size of 256, and the second linear layer maps these to the final output dimension corresponding to the number of classes.

The forward pass of the model performs the following operations:

1. Embed the input sequence.
2. Process the embeddings through the LSTM layer to obtain the last hidden state.
3. Concatenate the embeddings with the LSTM's last hidden state.
4. Apply a linear transformation followed by a ReLU activation.
5. Perform global max pooling on the resulting tensor.
6. Apply dropout for regularization.
7. Pass the pooled features through the final linear layer to obtain the output classification scores.

#### Mhcnuggets Prediction Hyperparameters

```
from mhcnuggets.src.predict import predict
predict(class_='I',
             peptides_path=peptides_path,
             mhc='{}'.format(allele), output='{}.txt'.format(allele))
```

This code snippet uses the MHCnuggets library to predict peptide bindings to MHC molecules. Specifically:

1. Import the prediction function from MHCnuggets.
2. Call the predict function, specifying:

- class_='I' for predicting bindings to MHC class I molecules (use 'II' for MHC class II).
- peptides_path as the path to a text file containing peptide sequences.
- mhc, set to a specific allele name to use for prediction.
- output, set to the filename where the prediction results will be saved, named after the allele.
- Each allele is processed individually in a loop, with the prediction results for each saved to a separate file.

#### Mhcflurry Prediction Hyperparameters

```
import mhcflurry
predictor = mhcflurry.Class1PresentationPredictor.load()
results = predictor.predict(peptides, alleles=alleles_list)
```

This code snippet demonstrates the use of the MHCflurry library in Python to predict the binding of peptide sequences to MHC class I molecules. Here's a brief overview:

- Import the MHCflurry library.
- Load a class I presentation predictor.
- Use the predict method to predict the binding between a list of peptides (peptides) and a list of MHC class I alleles (alleles_list).

Due to a limitation in MHCflurry, it can only process up to 6 alleles at a time. Therefore, a loop is typically used to handle all alleles by processing them in batches of six.

#### NetMHCpan Prediction Hyperparameters

```
../netMHCpan -p NEPDB_I_peptide.pep -BA -xls -xlsfile ../HLA_I_result/output_HLA-A01109.csv -a allele
```

This code snippet demonstrates the use of the NetMHCpan to predict the binding of peptide sequences to MHC alleles. Here's a brief overview:

- -p means the input consists of peptides.
- -BA specifies that the prediction mode is BA (Binding Affinity), which is more robust.
- -xls -xlsfile means the output is in XLS format.
- output_path specifies the path for the output files.
- -a specifies the allele name.

#### MixMHCpred Prediction Hyperparameters

```
MixMHCpred -i INPUT_FILE -o OUTPUT_FILE [-a LIST_OF_ALLELES]  [-p PEPTIDES_SCORING] [-m OUTPUT_MOTIFS]
```

This code snippet demonstrates the use of the MixMHCpred to predict the binding of peptide sequences to MHC alleles. Here's a brief overview:

- -i, --input: Absolute or relative path to the input file (FASTA format or list of peptides).
- -o, --output: Name of the output file or directory.
- -p, --peptides_scoring: Enable (1) or disable (0) binding predictions. Default is 1 for peptides scoring, 0 for sequence alignment.
- -m, --output_motifs: Enable (1) or disable (0) plotting of logos and creation of an HTML file for motifs. Default is 0.
- -a, --alleles: List of MHC alleles separated by commas

#### Anthem Prediction Hyperparameters

```
python sware_b_main.py --HLA "$allele" --mode prediction --peptide_file "$peptide_file"
```

This code snippet demonstrates the use of the Anthem to predict the binding of peptide sequences to MHC alleles. Here's a brief overview:

- --peptide_file: the path of the file that contains peptide sequence in text format or protein sequence in fasta format, respectively.
- --HLA: List of MHC alleles separated by commas
- --mode: prediction the mode choose to use. In train model function, if users want to use their trained model, the mode is "useYourOwnModel"

#### TransPHLA Prediction Hyperparameters

```
python pHLAIformer.py --peptide_file "peptide.fasta" --HLA_file "./HLA/HLA_allele.fasta" --threshold 0.5
--cut_peptide False --cut_length 14 --output_dir "./results/" --output_attention False --output_heatmap False
--output_mutation False
```

This code snippet demonstrates the use of the TransPHLA to predict the binding of peptide sequences to MHC alleles. Here's a brief overview:

- peptide_file: type = str, help = the filename of the .fasta file contains peptides
- HLA_file: type = str, help = the filename of the .fasta file contains sequence
- threshold: type = float, default = 0.5, help = the threshold to define predicted binder, float from 0 - 1, the recommended value is 0.5
- cut_peptide: type = bool, default = True, help = Whether to split peptides larger than cut_length?
- cut_length: type = int, default = 9, help = if there is a peptide sequence length > 15, we will segment the peptide according the length you choose, from 8 - 15
- output_dir: type = str, help = The directory where the output results are stored.
- output_attention, type = bool, default = True, help = Output the mutual influence of peptide and HLA on the binding?
- output_heatmap: type = bool, default = True, help = Visualize the mutual influence of peptide and HLA on the binding?
- output_mutation: type = bool, default = True, help = Whether to perform mutations with better affinity for each sample?

#### DeepSeqPanII Prediction Hyperparameters

We Match Alleles and Models: Ensure that each allele is matched with the appropriate model, And process sequences: Process all required sequences using the specified model and alleles.

To use DeepSeqPanII with similar parameters, you can follow these steps:

- Clone DeepSeqPanII: Clone the DeepSeqPanII repository into the desired folder with the command:
- git clone https://github.com/pcpLiu/DeepSeqPanII.git
- Switch Directory: After cloning, navigate to the code_and_dataset directory:
- cd DeepSeqPanII/code_and_dataset
- Use the Command: Execute the following command to run DeepSeqPanII:

```
python deepseqpanII.py <model_path> <allele1_name> <allele2_name> <peptide_sequence>
```

This code snippet demonstrates the use of the DeepSeqPanII to predict the binding of peptide sequences to MHC alleles. Here's a brief overview:

- <model_path> is the path where the model is located.
- <allele1_name> is the name of the first allele.
- <allele2_name> is the name of the second allele.
- <peptide_sequence> is the peptide sequence to be processed.

#### NetMHCIIpan Prediction Hyperparameters

```
./netMHCIIpan -inptype 1 -f NEPDB.pep -BA -xls -xlsfile  output_path -a allele
```

This code snippet demonstrates the use of the NetMHCIIpan to predict the binding of peptide sequences to MHC alleles. Here's a brief overview:

- -inptype 1 means the input consists of peptides.
- -BA specifies that the prediction mode is BA (Binding Affinity), which is more robust.
- -xls -xlsfile means the output is in XLS format.
- output_path specifies the path for the output files.
- -a specifies the allele name.

#### MixMHC2pred Prediction Hyperparameters

```
MixMHCpred -i INPUT_FILE -o OUTPUT_FILE [-a LIST_OF_ALLELES]  [-p PEPTIDES_SCORING] [-m OUTPUT_MOTIFS]
```

This code snippet demonstrates the use of the MixMHC2pred to predict the binding of peptide sequences to MHC alleles. Here's a brief overview:

- -i, --input: Absolute or relative path to the input file (FASTA format or list of peptides).
- -o, --output: Name of the output file or directory.
- -a, --alleles: List of MHC alleles separated by commas
- --no_context: Use the --no_context option in MixMHC2pred for analyzing pre-cleaved peptides without context, which is suitable for experiments testing specific peptides directly.
